# Assessing variation in *Citrus limon* ‘Assam lemon’: morphological, seed availability pattern, and biochemical diversity across different districts of Assam

**DOI:** 10.1101/2023.09.01.555945

**Authors:** Suraiya Akhtar, Raja Ahmed, Khaleda Begum, Ankur Das, Sarat Saikia, Sofia Banu

**Author notes:** Corresponding Author Email ID.

## Abstract

The Assam lemon is a highly valued *Citrus* cultivar known for its unique aroma, flavor, and seedless fruit characteristic. This study aimed to investigate for the first time the morphological, seeding pattern, and biochemical, variations within 133 populations of Assam lemon cultivar growing across 22 districts of Assam, India, and including control population, with the objective to offer comprehensive understanding that could facilitate the improvement of breeding programs and further improvement of this important cultivar. Using the dendrogram-based UPGMA algorithm, morphological and seeding pattern data were analyzed at both district and population levels. The resulting dendrograms revealed two major clusters, where all the populations of Upper Assam districts were in the same cluster with the original stock (control population). However, populations from Tinsukia and Dhemaji districts displayed more close similarities with the control population in comparison to other populations of Upper Assam districts. Another interesting observation was regarding flowering patterns, while some populations displayed both bisexual and unisexual flowers with less concentration of unisexual flowers, others had bisexual and unisexual flowers of almost equal concentration. Unisexual flowers contained only the male reproductive organs with 40 anthers, while bisexual flowers had 36 anthers. Seeding patterns were examined across the districts, and it was found that populations from Tinsukia, Dhemaji, Lakhimpur, Dibrugarh, Jorhat, and the control population exhibited seedless characteristic while populations from other selected districts displayed a combination of seedless and seeded traits. Interestingly, Golaghat district appears as the linking district and showed availability of both seeded and seedless Assam lemon fruit, connecting the regions of Upper, Central, Lower and North Assam and Barak valley. Biochemical analysis showed significant variations across districts, however, the populations from Dhemaji, Tinsukia, Lakhimpur, Dibrugarh, and Jorhat districts displayed similarity with the control population. The study also investigated variability in soil nutrient content revealing substantial variation among the populations studied, potentially influencing observed morphological variations. This comprehensive investigation provides valuable insights into the morphological, seeding pattern, and biochemical diversity within the Assam lemon cultivar. These findings can be instrumental in breeding programs to enhance this *Citrus* cultivar, particularly in producing high-quality seedless fruits to meet consumer demands.

## Introduction

*Citrus* fruits are globally renowned for their economic importance, nutritional value, and cultural significance (Saeid and Ahmed 2021). In India, Assam stands as a prominent region for *Citrus* cultivation, encompassing diverse *Citrus* species with district flavours, aromas, and appearance (Hore and Barua 2004). Among this, Assam lemon, an important cultivar of *C. limon* holds a special place as a widely grown *Citrus* cultivar in Assam (Sheikh et al. 2021). Known for its distinct aroma, and flavour, Assam lemon has gained popularity in both domestic and international markets (Ahmed et al. 2023). Further, due to its nutritional value, it is a great source for the manufacture of several medicinal and cosmetic products (Barua and Bharadwaj 2017). The morphology of Assam lemon exhibits a range of traits, including fruit size, fruit shape, fruit colour, peel texture, seed numbers etc. (CNTUS 2019). These morphological variations may be influenced by factors such as soil type, rainfall, and agricultural practices followed by farmers in different districts (Liliane and Charles 2019). The understanding of morphological variation within Assam lemon cultivar across different districts of Assam is expected to assist in breeding programs for further improvement of Assam lemon (Jain et al. 2017).

Assam lemon forayed into the cultivation and market after being identified at by Government *Citrus* Fruit Research Station, Byrnihat (Thakur 2010). In the 1940s, S.C. Bhattacharyya and S. Dutta, Horticulture Development Officer and Officer-in-charge respectively, of Government *Citrus* Fruit Research Station (GCFRS), Byrnihat, collected a variety from Hahchora village under the name “China-kaghi” (Ahmed et al. 2023). While experimenting with the seeds, they stumbled upon a seedling which was characteristically different from others and had seedless fruit (Thakur 2010). In order to preserve this unique quality of non-seediness, it was further propagated by vegetative means and developed as a clonal horticultural cultivar and named as Assam lemon as documented by the in-monograph Classification of *Citrus* Fruits of Assam published by the Indian Council of Agricultural Research in 1956 (Bhattacharya and Dutta 1956). Assam lemon, a lemon like no other, boast an extraordinary aroma, flavour, and seedlessness that set it apart from its counterparts (Thakur 2010). Farmers, both commercial and domestic, have embraced the lemon’s unique qualities, cultivating it in homestead gardens as a standalone crop or alongside other *Citrus* varieties (Thakur 2010). *Citrus* species, including Assam lemon, have a tendency towards natural hybridization, which makes it highly labile to acquire variations in seed formation in different populations over time (Velasco and Licciardello 2014, Wu et al. 2014). Moreover, study by Kahn et al. (2004) highlighted that specific *Citrus* varieties can yield seedless fruits due to limited functional pollen and ovules, regardless of their neighboring varieties. Conversely, some *Citrus* types may possess ample functional pollens and ovules but are self-incompatible, resulting in seedless fruits when grown alone. However, when grown near cross-pollinating varieties, these self-incompatible types can produce seedy fruits (Kahn et al. 2004). Assam lemon has acquired a prized status as the second most grown *Citrus* variety of Assam with about more than 15 thousand hectares of cultivated area and an annual production of more than 1.56 lakh metric tons (Government of Assam Database 2022). While, the genetic diversity of Assam lemon within different districts of Assam has been established through ISSR marker analysis, the seeding pattern in Assam lemon fruits has not been extensively studied across different regions of Assam (Ahmed et al. 2023). Seedlessness is an important character because of the consumer preference for seedless fruits, which offer convenience and culinary applications (Premachandran et al. 2019) and Assam lemon has gained popularity over the last few decades because of these desired features. However, recent survey of available market fruits has shown existence of both seeded and non-seeded fruits of Assam lemon, which emphasizes the need to study the seeding pattern among the fruits being grown across Assam. This information can guide *Citrus* growers in making informed decisions regarding the most suitable accessions for Assam lemon cultivation in different regions of Northeast India (Emery and Offord 2019).

In addition to morphological and seeding variations, the biochemical composition is known to play a crucial role in determining its nutritional value and potential health benefits of Assam lemon (Samtiya et al. 2021). Recent investigations show that Assam lemon is rich several important compounds including citric acid, ascorbic acid, pectin, flavonoids etc. (Gogoi et al. 2023) that contribute to the antioxidant properties and other therapeutic attributes (Gogoi et al. 2023; Khatiwora et al. 2017). However, the concentration of these bioactive compounds may vary among the accessions from different districts of Assam owing to different environmental factors and agricultural practices (Vita et al. 2018). Investigating the biochemical variations in Assam lemon across different districts of Assam is expected to provide valuable information for understanding its potential of different populations for applications in functional foods, nutraceuticals, and pharmaceutical industry (Hsouna et al. 2023; Ye et al. 2016).

Variations in soil composition, including factors such as pH, and the micronutrient and macronutrients quantity may influence the growth, development, and productivity of Assam lemon (Morgan and Connolly 2013). By studying the variation in different soil parameters across districts of Assam, a better understanding of soil-Assam lemon interactions can be gained (Smith et al. 2022). This knowledge is valuable for optimizing soil management practices, including fertilizer application, irrigation, and soil amendment strategies, to enhance the productivity and quality of Assam lemon (Silver et al. 2021).

This study thus aims to provide a basic and comprehensive understanding of the morphological variation, soil variation, seeding pattern, and biochemical variation of Assam lemon in different districts of Assam. The findings from this study will benefit *Citrus* growers, breeders, and researchers involved in the cultivation and promotion of Assam lemon.

## Materials and Method

### Study Area Selection

In order to assess the diversity in morphological, seeding pattern, and biochemical characteristics of Assam lemon, a comprehensive study was conducted across all 22 districts of Assam (Figure 1). The selected districts from Upper Assam include Dhemaji, Dibrugarh, Tinsukia, Lakhimpur, Jorhat, and Golaghat; from Central Assam, the included districts are Karbi Anglong, Dima Hasao, Nagaon, and Morigaon; the districts from North Assam includes Udalguri, and Sonitpur; from Lower Assam, the districts included are Kamrup Metropolitan, Kamrup Rural, Baksha, Nalbari, Barpeta, Bongaigaon, Kokrajhar, and Dhubri; and the districts from Barak Valley included Cachar, and Karimganj (Figure 2-6). This careful selection ensured the representation of all regions within Assam for the purposes of this investigation. These districts represent a range of geographical populations, elevation, and soil types prevalent in Assam. Furthermore, fruits were also taken from the Horticulture Research Station, Kahikuchi to serve as a control population (Table SF 1).

**Figure 1:**
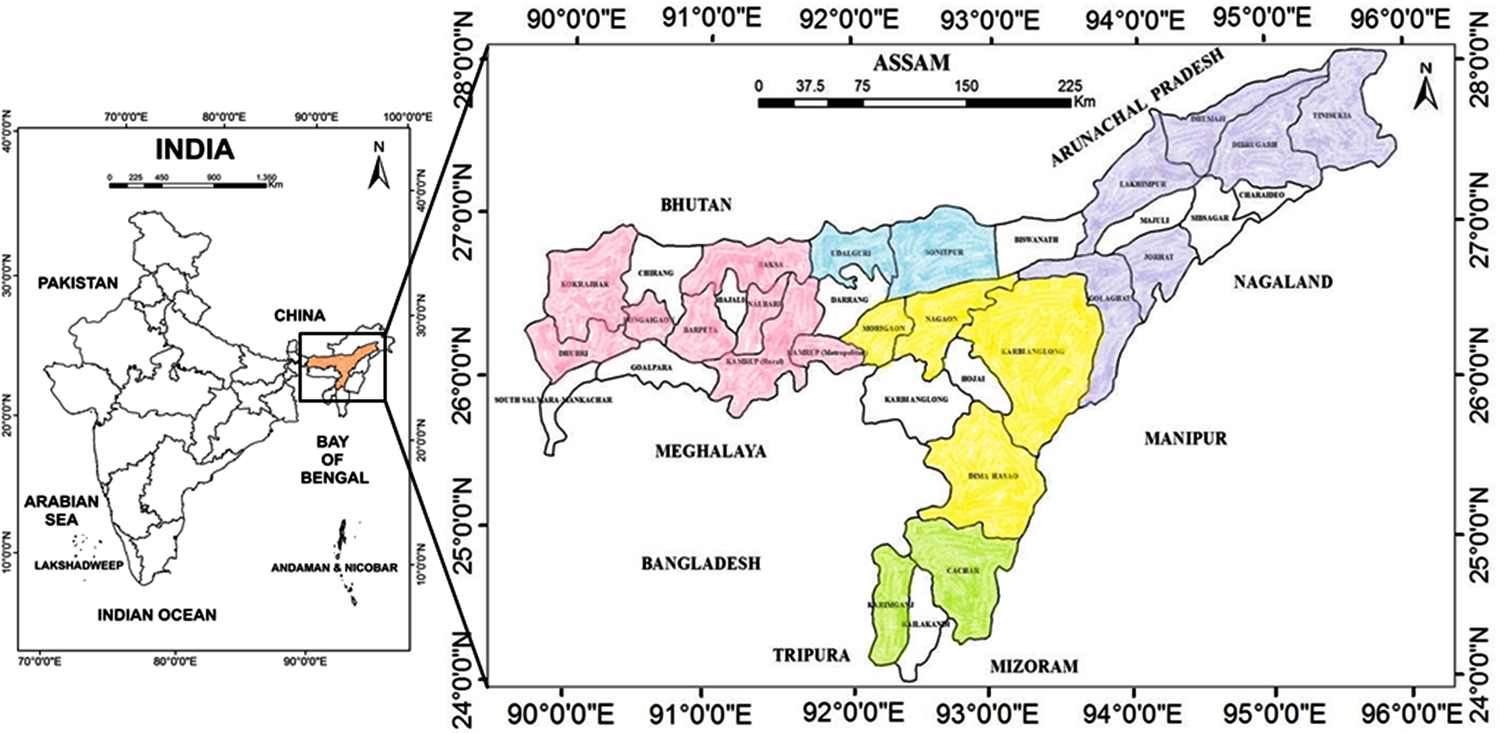
Map of Assam, India showing the study area

**Figure 2:**
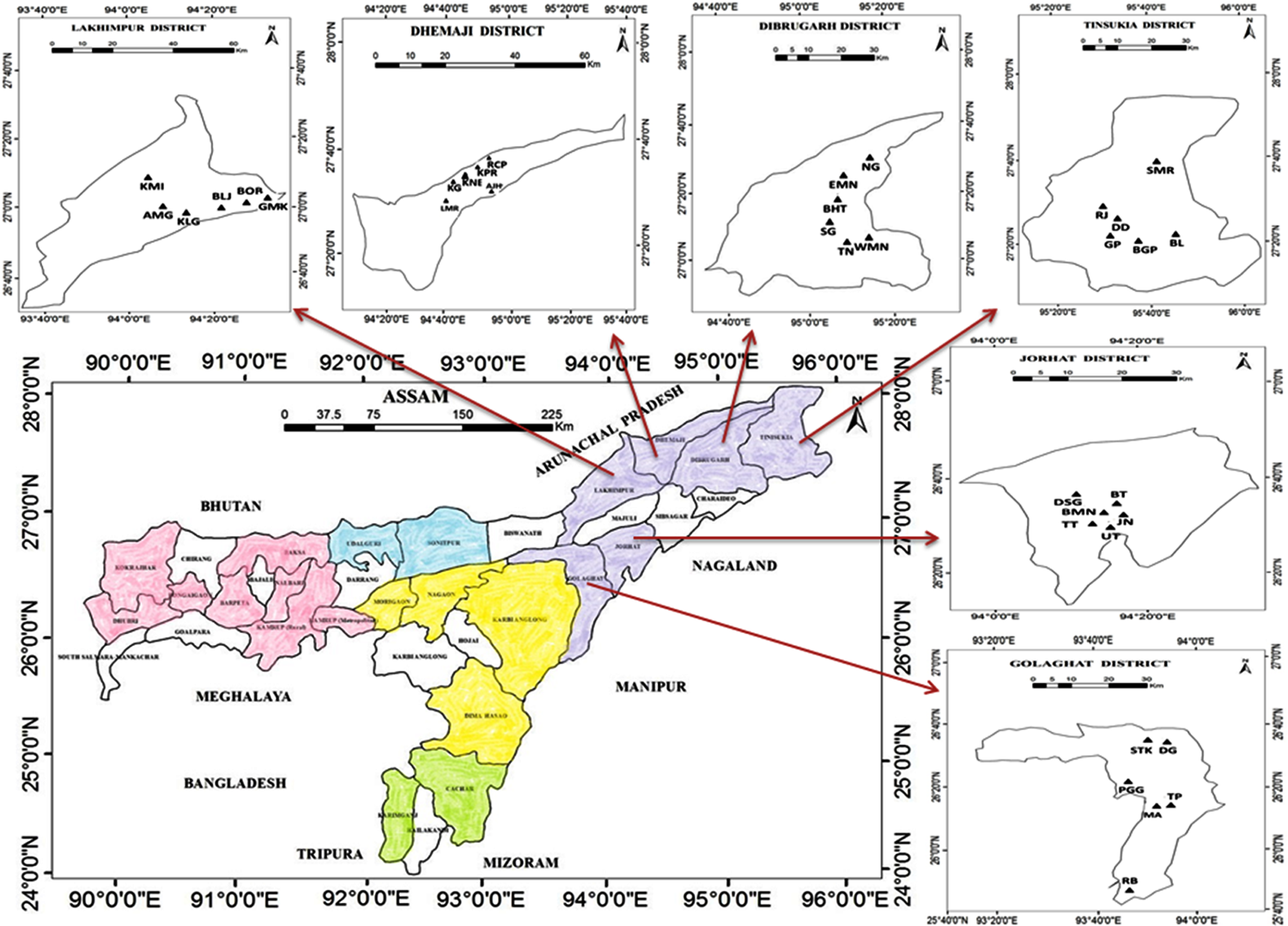
Map of Assam showing the geographic locations of the sampled population in Upper Assam

**Figure 3:**
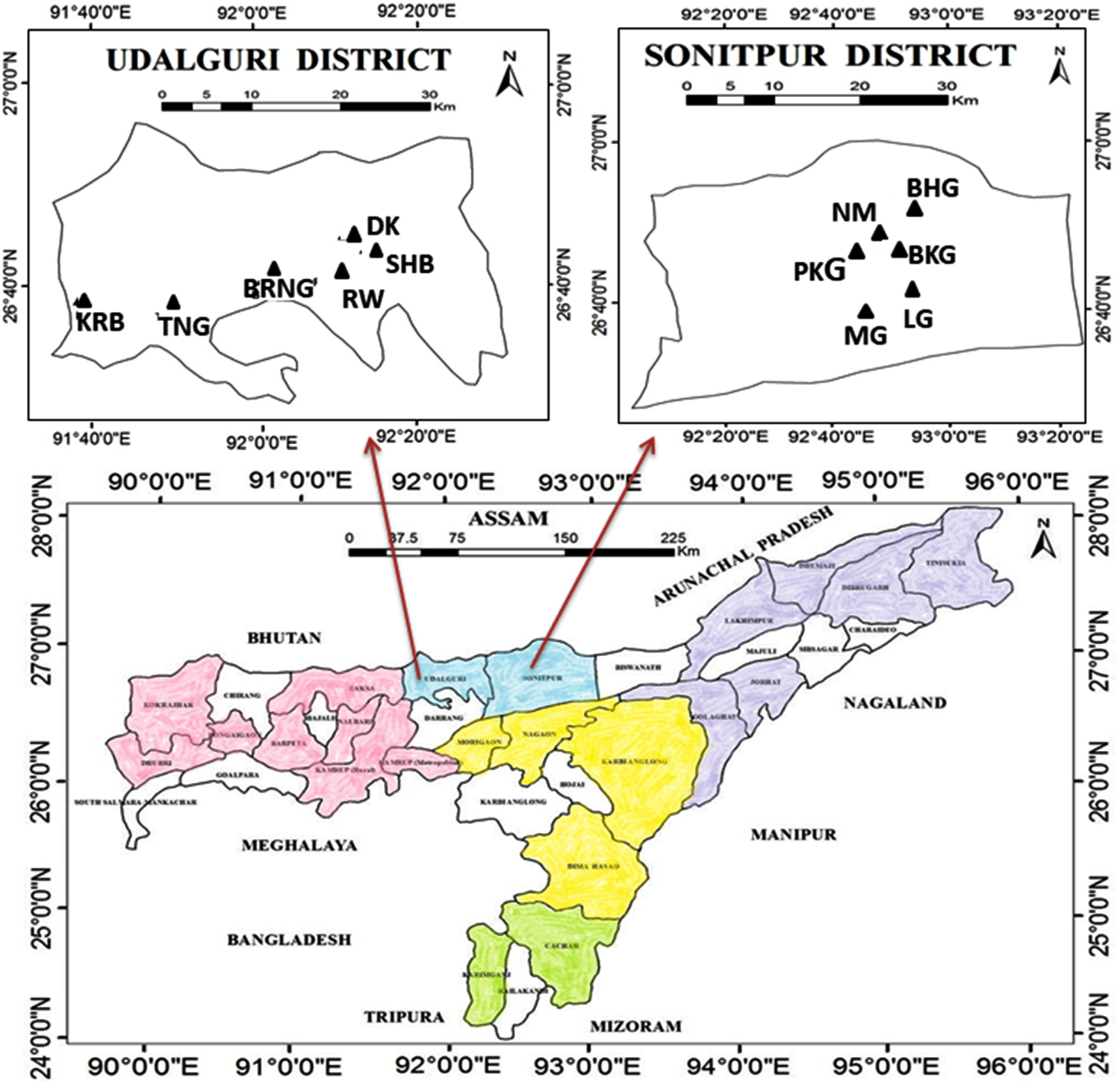
Map of Assam showing the geographic locations of the sampled population in North Assam

**Figure 4:**
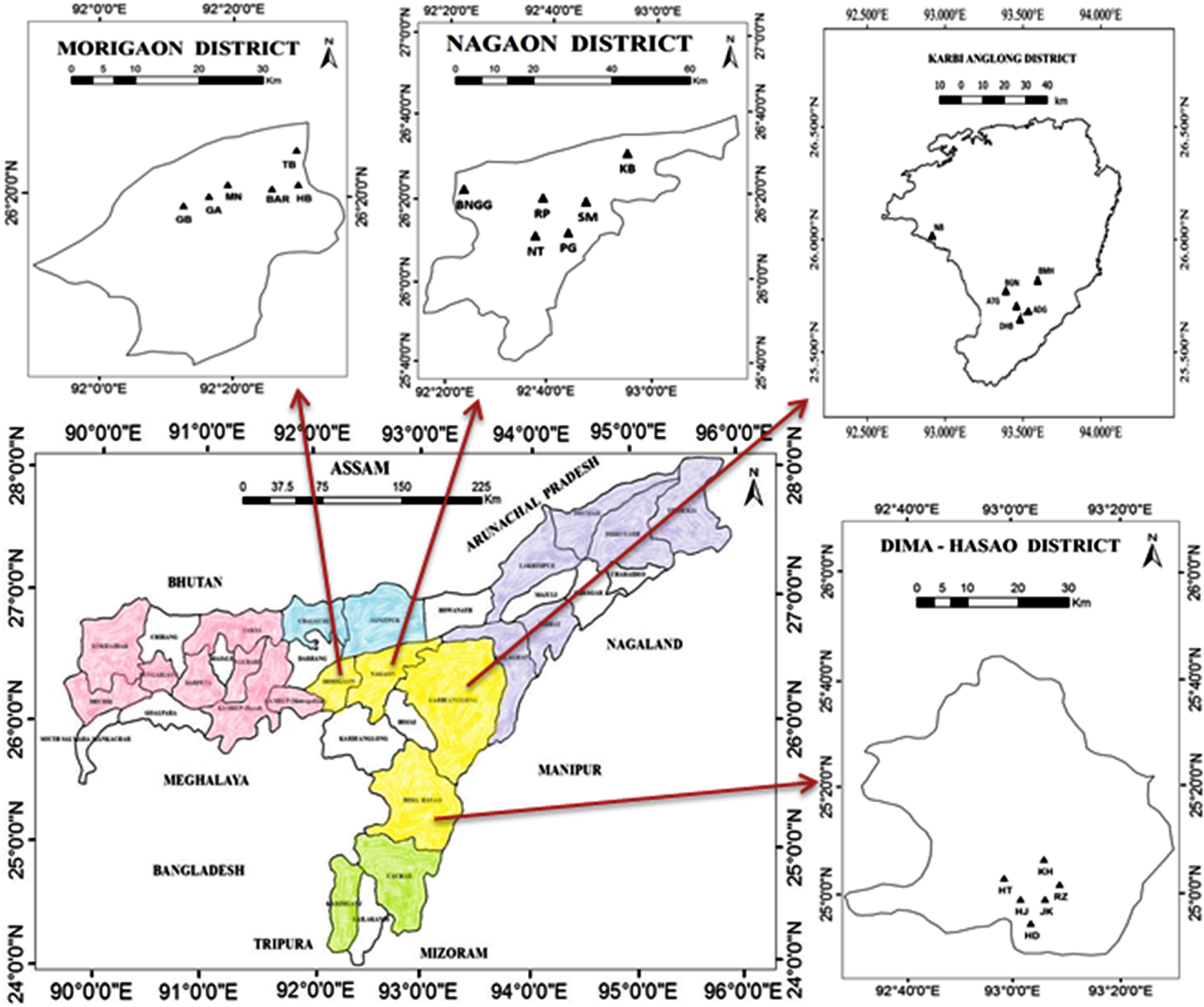
Map of Assam showing the geographic locations of the sampled population in Central Assam

**Figure 5:**
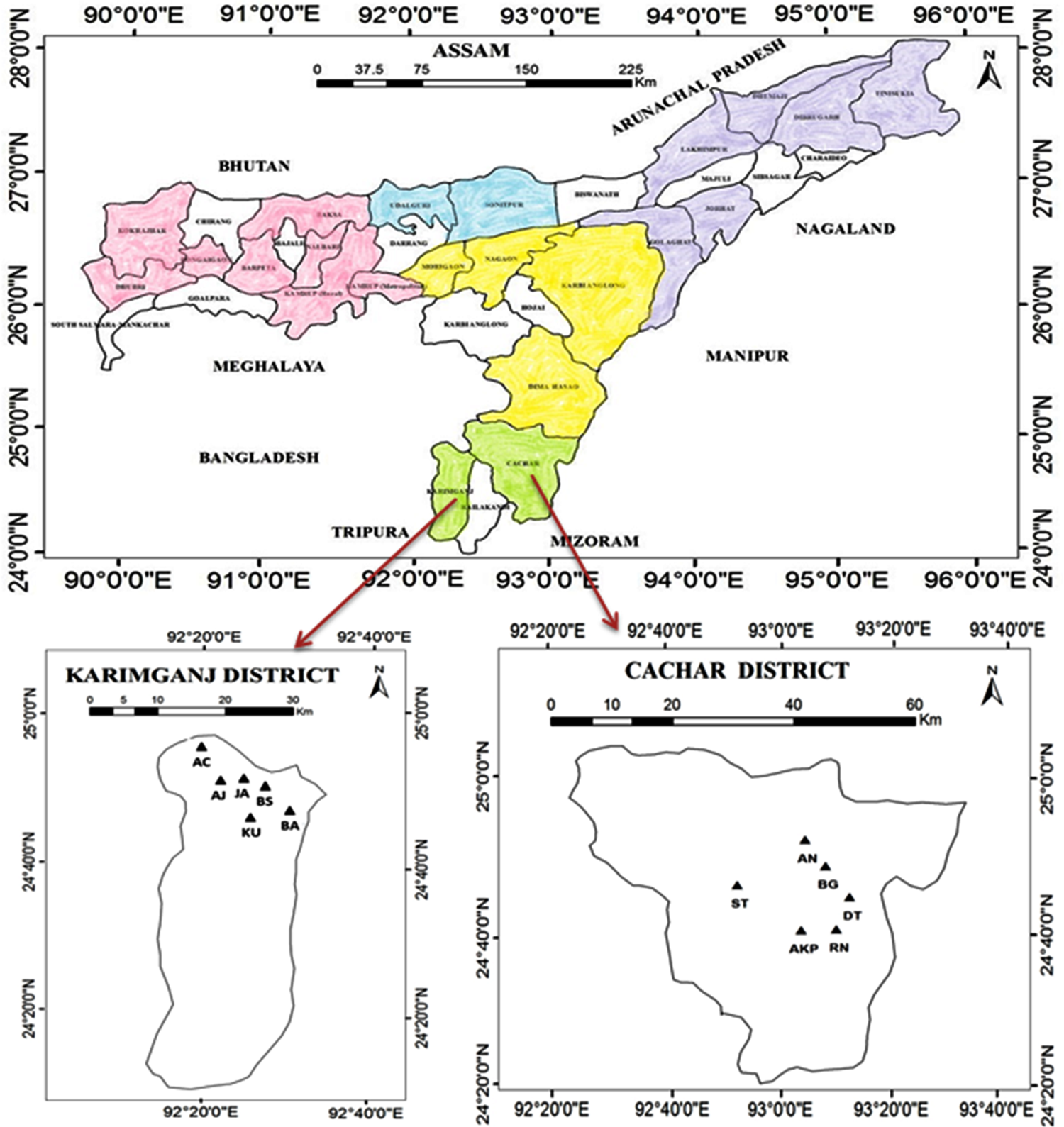
Map of Assam showing the geographic locations of the sampled population in Barak Valley

**Figure 6:**
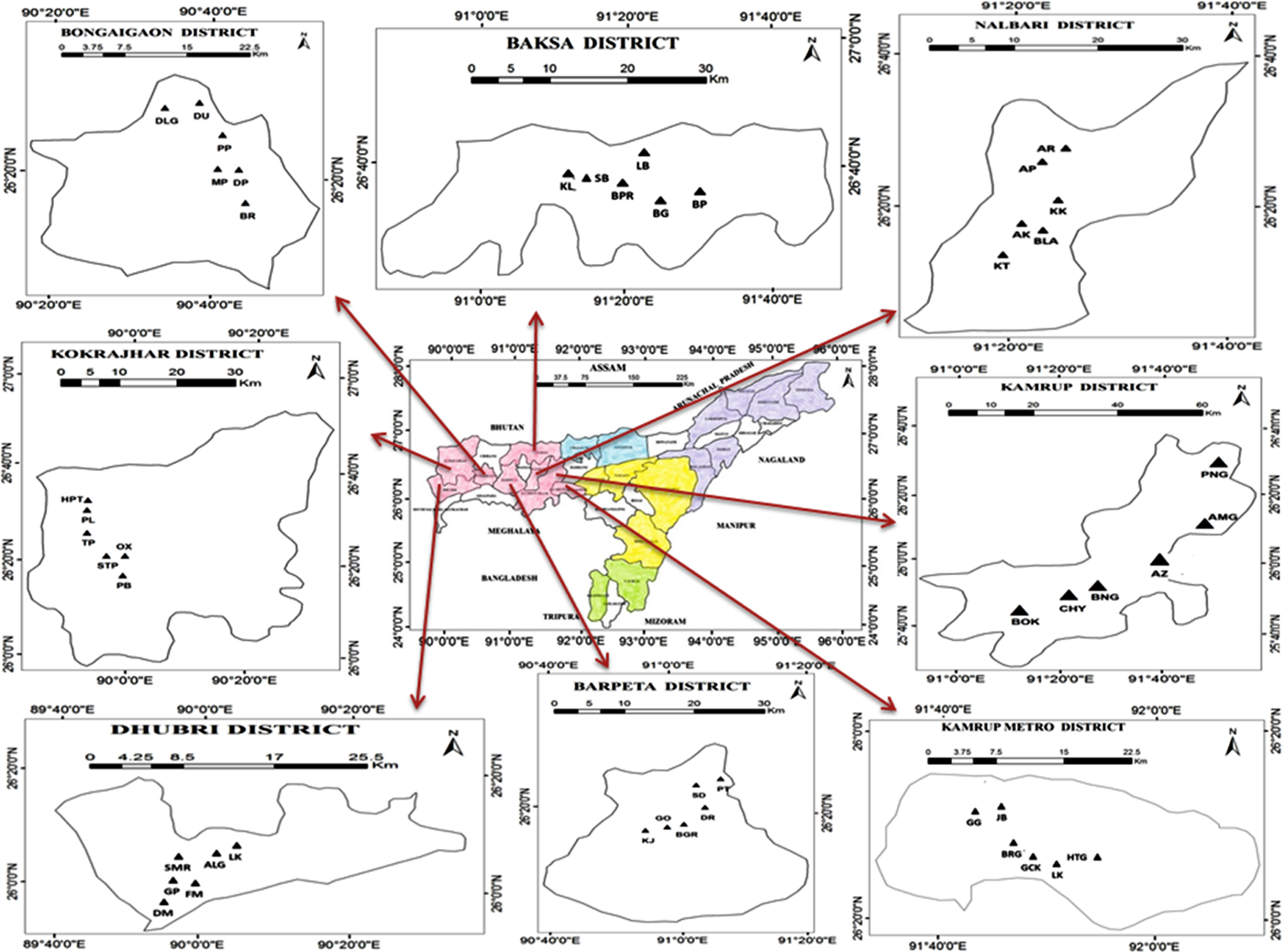
Map of Assam showing the geographic locations of the sampled population in Lower Assam

### Fruit Materials

Fresh fruit samples of Assam lemon were systematically collected from a total of 132 populations across 22 selected districts in Assam. Additionally, samples were also obtained from the Horticulture Research Station, Kahikuchi, which served as the control population for comparison. These collected fruit samples were carefully handled and stored in a refrigerated environment at −80°C until the completion of the present investigation (Table SF 1).

### Morphological Analysis

For our research on morphological diversity across the various districts of Assam, we carefully considered a comprehensive set of 59 variables. These variables encompass a wide range of characteristics related to trees, including their physical traits, leaves, flowers, fruits, seeds etc.

To ensure a thorough analysis, we examined the following aspects which includes, tree character, tree height, branches density, branches angle, thorn density, thorn size, thorn shape, leaf density, leaf division, leaf venation, leaf shape, leaf apex, leaf margin, leaf intensity of green, leaf size, leaf length, flower nature, flower sexuality, flower intensity, flower color, flower length (cm), sepal arrangement, sepal length (cm), petal arrangement, petal length (cm), androecium arrangement, androecium length (cm), number of anthers, anther length (cm), anther color, anther arrangement, filament length (cm), filament color, gynoecium arrangement, gynoecium length (cm), flowering shoot, fruit density, fruit color, fruit shape, fruit skin texture, fruit base, fruit apex, fruit axis, fruit weight, fruit length, fruit diameter, specific gravity, number of segments, fruit peel weight, fruit pulp weight, fruit pulp: peel ratio, fruit rind thickness (mm), albedo thickness (mm), flavedo thickness (mm), the length from pith to albedo (mm), pith diameter (mm), seed availability, seed number, seed color. The data collected was further analyzed using statistical techniques and clustering methods, to identify patterns, relationships, and potentially distinct groups of Assam lemon within the studied regions.

### Seeding Pattern

For the investigation of seeding pattern of Assam lemon, mature fruits were sampled from different populations of 22 districts throughout the Assam. In addition, fruits were collected from the Horticulture Research Station, Kahikuchi to serve as a control population for comparison. The mature fruits were carefully collected at their maturity stage (60 DAF) to ensure accurate seed analysis. In the laboratory, the collected fruits were gently washed and dried to remove any external contaminants. Subsequently, each fruit was cut open, and the seeds were carefully extracted using a sterile knife, and forceps. The total number of seeds per fruit was counted, and their color was recorded. The data obtained from the seed analysis were subjected to statistical analysis using methods described below to determine the average number of seeds per fruit and assess any significant variations among different samples (Table SF 1).

### Biochemical Analysis

The fruit quality characteristics of selected districts of Assam along with control population were assessed, focusing on several parameters (Table SF 1). These parameters included pH level, % of juice content, total soluble solids (measured in °Brix), citric acid concentration (measured in g/ml), ascorbic acid concentration (measured in mg/ml), total soluble solids to titratable acidity ratio, total sugar content (measured in µg/ml), reducing sugar content (measured in µg/ml), carotenoid concentration (measured in mg/g), chlorophyll content (measured in µg/g), pectin content (both in the peel and pulp), equivalent weight (for both peel and pulp), methoxy content (for both peel and pulp), anhydrounic acid content (for both peel and pulp), and degree of esterification (for both peel and pulp).

### pH

Total pH of the Assam lemon juice were measured using pH 700 meter (Eutech Instruments, United Kingdom) (Reddy et al. 2016).

### Total Soluble Solids

Total soluble solids of the lemon juice were determined as °Brix using OPTi digital refractometer (Bellingham + Stanley, United Kingdom) (Uresti-Porras et al. 2021).

### % Juice Content

% juice content determination was performed following the methodology described by Kashyap et al. in 2020, with slight modifications. The weight of both the fruit and its corresponding juice content were measured in grams and documented. The % of juice content was then calculated using the formula given below,

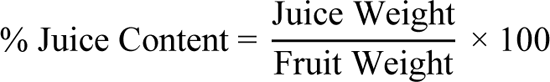

### Citric Acid

The determination of citric acid content in the juice of Assam lemon fruits was done using a modified protocol described by Brima et al. in 2014. To measure the concentration of citric acid, the lemon juice was extracted by squeezing the fruits and then diluted with distilled water at a ratio of 1:9 (Juice:Water). Phenolphthalein was added as an indicator, and the titration was conducted using NaOH as the base. The titration process continued until the color of the juice turned red/pink and remained consistent for at least 15 seconds. This titration was repeated with additional aliquots of the sample solution until concordant results were obtained. The concentration of citric acid was then calculated using the formula given below,

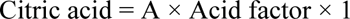

Where,

A = Mean value of coordinate readings Acid factor = 0.0064

### Ascorbic Acid

The determination of ascorbic acid content in the juice of Assam lemon fruits, a modified protocol described by Satpathy et al. in 2021 was followed. To measure the concentration of ascorbic acid, 20 ml of fresh lemon juice filtrate was transferred into a 250 ml conical flask, and 1 ml of starch indicator was added. The sample solution was then titrated against a 0.01 mol L^-1^ iodine solution until the color changed to a dark black color. The titration process was repeated with additional aliquots of the sample solution until concordant results were obtained. The concentration of ascorbic acid was then calculated using the formula given below,

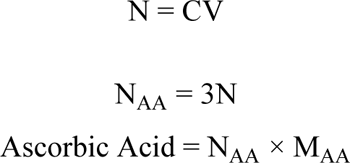

Where,

N = Moles of iodine

C = 0.01mol L^-1^

V = Mean value of the conordant readings

N_AA_ = Moles of ascorbic acid

M_AA_ = Molecular mass of ascorbic acid

### Total soluble solids/Titratable Acidity (TSS/TA)

The TSS/TA ratio of Assam lemon fruit was assessed using the methodology outlined in the study conducted by Kashyap et al. in 2020. This ratio was obtained by dividing the °Brix value (representing the Total Soluble Solids) by the percentage of acid content.

### Total Sugar and Reducing Sugar

The quantification of reducing sugar was performed using 3,5-dinitro salicylic acid (DNS), following the modified protocol described by Gusakov et al. in 2011. Additionally, the determination of total sugar was conducted using Anthrone reagent, as outlined by Buckan in 2015 with slight modifications.

### Carotenoid Content

The carotenoid content of the peel samples from Assam lemon was determined using the methodology described by Kashyap et al. in 2020, with slight modifications. 1 g sample of fruit was ground with liquid nitrogen, and the resulting powder was mixed with 10 ml of a hexane:acetone:ethanol solution (v/v; 50:25:25). Subsequently, the mixture was centrifuged at 4000 g for 5 minutes. The supernatant, containing the color compounds, was collected, and the volume was adjusted to 10 ml with the extraction solvent. The total carotenoid content was then measured by assessing the absorbance at 450 nm using a spectrophotometer. The concentration of carotenoids was calculated using the following equation,

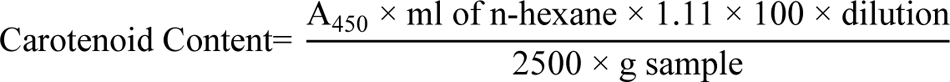

Where,

A= Absorbance at 450 nm wavelength

V= Total volume of sample

W= Weight of fresh plant tissue

### Chlorophyll Content

The chlorophyll content of the juice of Assam lemon at various fruit developmental stages were determined using the protocol described by Kashyap et al. 2020 with minor modification. For the determination of chlorophyll concentration in the fruit peel, 1 g of peel sample was ground with liquid nitrogen. The resulting powder was then mixed with 20 ml of 80% acetone solution containing 0.5 g of MgCO_3_. After incubating the mixture at 4°C for 3 hours, it was centrifuged at 2500 rpm for 5 minutes. The supernatant was carefully collected, and its absorbance was measured at 654 nm and 663 nm using a spectrophotometer. Using the provided equation, the concentration of chlorophyll was calculated.

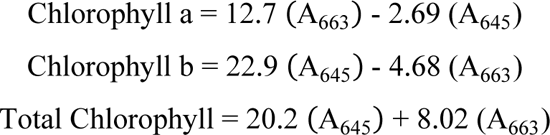

Where,

A= Absorbance at specific wavelength

V= Final volume of chlorophyll extract in 80% acetone

W= Fresh weight of tissue extracted

### Pectin Content

The determination of pectin content in both the peel and pulp samples of Assam lemon was done following the protocol outlined by Khamsucharit et al. in 2017, with minor modifications. A total of 5 g of dried fruit sample was mixed with 100 ml of Citric acid solution (prepared by mixing 9 parts distilled water with 1 part Citric acid) and incubated at 65°C for 1 hour. After the completion of incubation, the extract was filtered, and 95% ethanol was added to the filtrate in a 1:1 proportion. The mixture was then incubated at room temperature for 2 hours. Following

incubation, the flocculants were skimmed off and washed 2-3 times using ethyl alcohol. The resulting precipitate was thoroughly dried at 35°C to 40°C and weighed to determine the pectin concentration. The concentration of pectin was calculated using the provided equation.

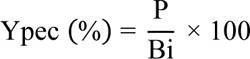

Where,

Ypec = Yield of pectin

P = Amount of extracted pectin

Bi = Initial amount of fruit powder

### Equivalent Weight

The determination of equivalent weight content in both the peel and pulp samples of Assam lemon was done following the protocol outlined by Khamsucharit et al. in 2017, with minor modifications. In this process, 0.5 g of dried pectin sample obtained from the previous pectin estimation experiment was dissolved in 5 ml of ethanol. To this pectin solution, 1 g of NaCl and a few drops of phenol red indicator were added. The resulting mixture was titrated against 0.1 M NaOH until a pale pink color, indicating the end point, was achieved. The titration was repeated with additional aliquots of the sample solution until consistent and concordant results were obtained. The equivalent weight was then calculated using the provided equation.

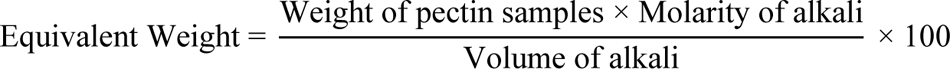

### Methoxy Content (MeO)

The determination of methoxy content in both the peel and pulp samples of Assam lemon was done following the protocol outlined by Khamsucharit et al. in 2017, with minor modifications. The methoxy content was estimated using the neutralized titrated solution obtained from the equivalent weight estimation. To the solution, 0.25 M NaOH was added and stirred for 30 minutes, followed by the addition of 25 ml of 0.25 N HCl. The resulting mixture was titrated against 0.1 N NaOH until a pale pink color, indicating the end point, was achieved. The titration was repeated with additional aliquots of the sample solution until consistent and concordant results were obtained. The methoxy content was then calculated using the provided equation.

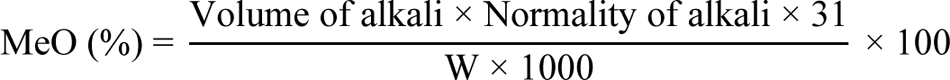

### Anhydrounic Acid (AUA)

The anhydrounic acid content in both the peel and pulp samples of Assam lemon was determined according to the method described by Khamsucharit et al. in 2017, with slight modifications. The calculation of anhydrounic acid content was performed using the provided equation.

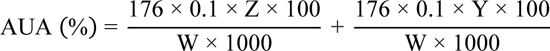

Where,

Molecular weight = 176 g

Z = Equivalent weight titration result

Y = Methoxy content titration result

W = Weight of sample

### Degree of Esterification (DE)

The degree of esterification in both the peel and pulp samples of Assam lemon was determined using the method described by Khamsucharit et al. in 2017, with slight modifications. The calculation of the degree of esterification was carried out using the equation provided.

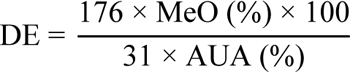

Where,

DE > 50% = High methoxy pectins (HM)

DE < 50% = Low methoxy pectins (LM)

### Statistical Analysis

For statistical analysis, the numerical data was generated using morphological characters including seeding pattern and also biochemical attributes which was used for the statistical analysis during the current investigation (Bardel-Kahr and Schliep 2023; Alaida and Aldhebiani 2022). Further, using the PAST 4.11 program, we performed, Principal Coordinate Analysis (PCoA), and constructed a dendrogram using the Unweighted Pair Group Method with Arithmetic Mean (UPGMA) based on these numerical data generated using morphological characters. Also, using the PAST 4.11 program, Analysis of Variance (ANOVA) test was performed using morphological and biochemical attributes.

### Soil Nutrient Estimation

Soil samples were carefully collected from various districts of Assam, focusing on locations with thriving Assam lemon trees during their active growth phase. Care was taken to avoid areas with apparent disturbances. Soil samples were collected to a depth of 15 centimeters, and several sub-samples were combined to create a composite sample at each site (TNAU Agritech Portal 2013). The collected soil samples were preserved and transported to the laboratory for analysis, where various physicochemical properties, including, pH, availability of macronutrients and micronutrients were measured. To assess the soil conditions in the 22 districts of Assam along with the control population, we employed soil testing kits (K054 - 1KT and K095L - 1KT, Himedia) (Table SF 1). These kits enabled us to estimate the concentrations of both micronutrients and macronutrients present in the soil samples collected for the current investigation. In terms of micronutrients, we focused on assessing the levels of Copper, Zinc, Boron, Manganese, Iron, and Molybdenum. These elements play crucial roles in plant growth and development, and their availability in the soil can significantly impact the health and productivity of vegetation. For macronutrient analysis, we evaluated several key parameters, including pH levels, organic Carbon content, Phosphate concentration, Potassium levels, Ammoniacal Nitrogen content, and Nitrate Nitrogen levels. These macronutrients are essential for plant nutrition, as they are involved in various physiological processes and influence overall soil fertility.

## Results

### Morphological Analysis

The analysis of morphological characters in Assam lemon revealed significant variation across different regions of the state (Table SF 2, Table SF 3, Table SF 4). The study encompassed a total of 59 morphological characters of various aspects, including tree characteristics, thorn characters, leaf traits, flower attributes, fruit properties, and seed features. Among all the studied characters, the flower traits exhibited the most pronounced variation. Notably, the investigation of Assam lemon flowers indicated that accessions from control, Dhemaji, Tinsukia, Jorhat, and Dibrugarh displayed both bisexual and unisexual flowers but, the concentration of unisexual flowers was comparatively quite less in these regions. In contrast, accessions from Golaghat, Central Assam (Karbi Anglong, Dima Hasao, Nagaon, and Morigaon), North Assam (Udalguri, and Sonitpur), Lower Assam (Kamrup Metropolitan, Kamrup Rural, Baksha, Nalbari, Barpeta, Bongaigaon, Kokrajhar, and Dhubri) to Barak valley (Dima Hasao, Cachar, and Karimganj) exhibited a combination of bisexual and unisexual flowers, having unisexual flowers of almost equal concentration to that of bisexual flowers. The unisexual flowers possessed only the androecium (Figure 7). Furthermore, the study revealed a distinct difference in the number of anthers between unisexual and bisexual flowers. Unisexual flowers were found to have as much as 40 anthers, whereas bisexual flowers exhibited 36 anthers (Table SF 3). Additionally, significant variations in fruit morphological characters were also observed across 690 accessions of Assam (Figure 8, Table SF 4).

**Figure 7:**
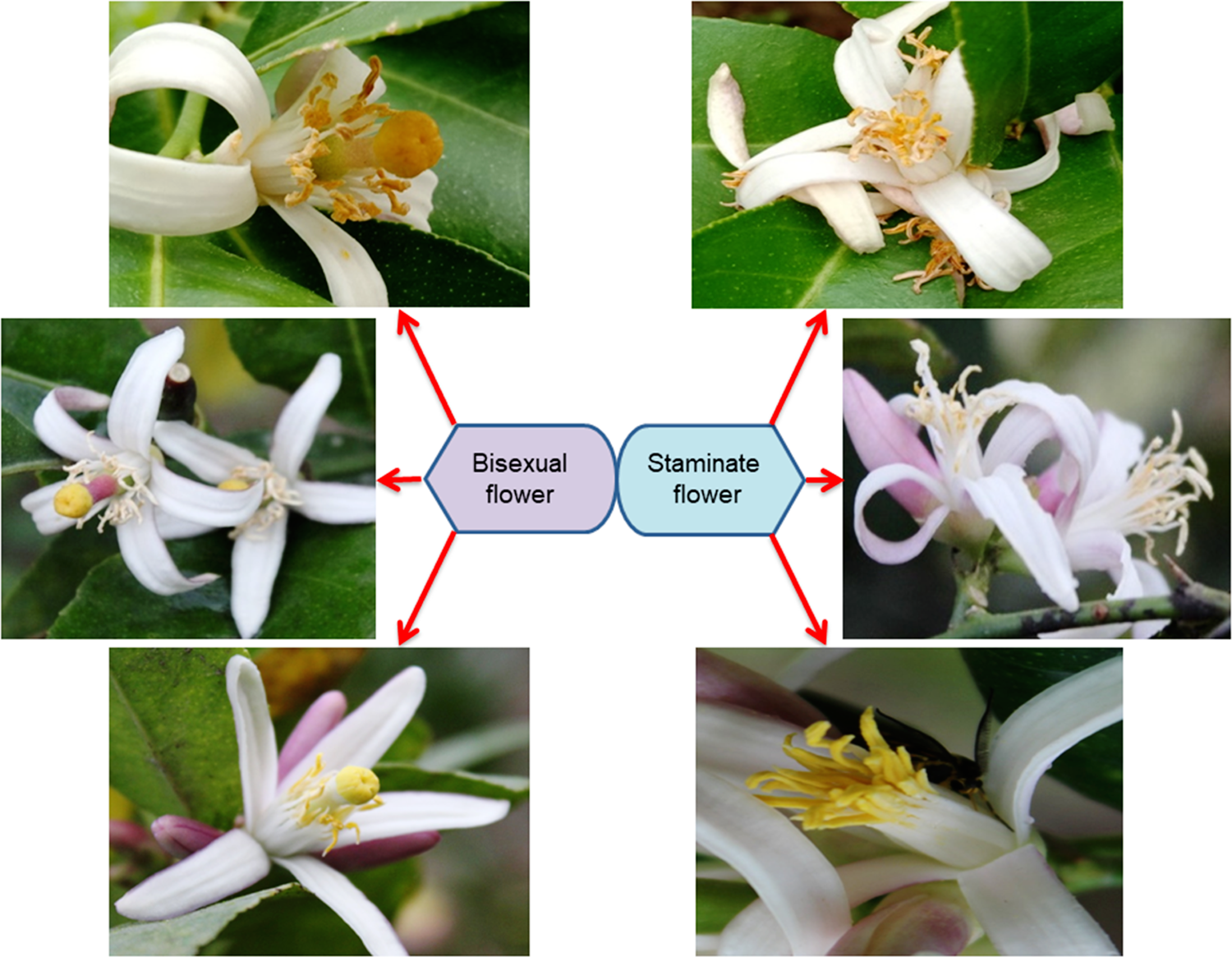
Floral morphology of Assam lemon: Bisexual flower (left); Staminate flower (right)

**Figure 8:**
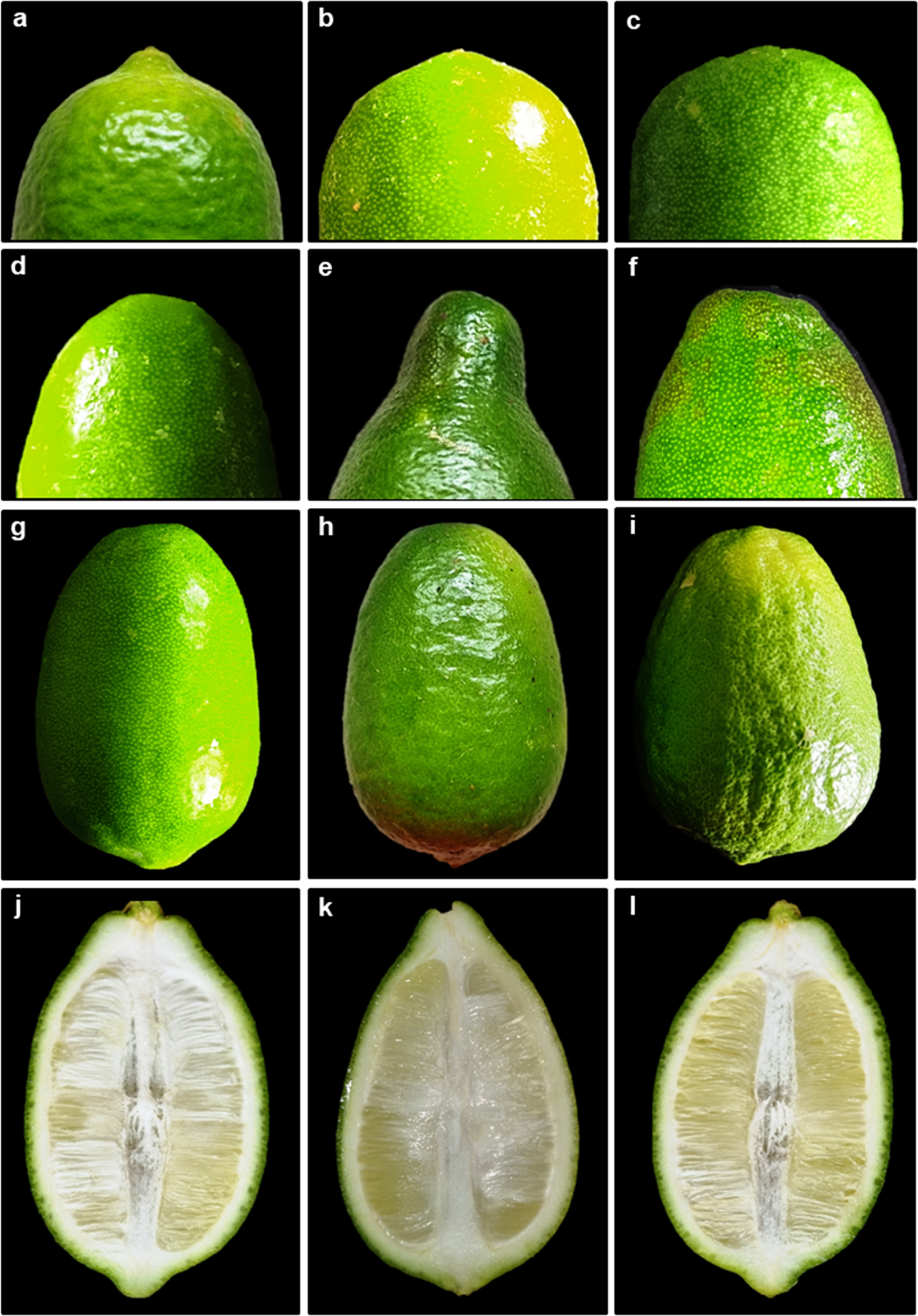
Fruit attributes of Assam lemon; **a-c:** Shape of the base of the fruit (**a:** mammiform, **b:** acute, **c:** round), **d-f:** shape of the apex of the fruit (**d:** round, **e:** necked, **f:** truncate), **g-i:** texture of the fruit (**g:** smooth, **h:** rough, **i:** bumpy), **j-l:** fruit axis (**j:** semi-hollow, **k:** solid, **l:** hollow)

### Seeding Pattern

During the investigation into the seeding pattern of Assam lemon across the 22 districts of Assam, an interesting finding emerged. It was observed that there is a combination of seeding patterns within the studied districts, with some districts exhibiting the true seedless trait characteristic of Assam lemon, while others exhibited a mixed character and had both seedless and seeded type (Figure 9). Also, we have observed that in the study area of Upper Assam districts and control, the Assam lemon trees were cultivated as a standalone crop. However, in other selected districts, the Assam lemon trees were found to be cultivated as standalone or alongside other *Citrus* varieties. Throughout the current investigation, we observed that the fruits collected from Tinsukia, Dhemaji, Lakhimpur, Dibrugarh, and Jorhat districts including control population were seedless. However, during the current investigation, we found a mixture of seedless and seeded fruits in the populations of Golaghat district, districts of Central Assam, North Assam, Lower Assam and Barak Valley districts. Interestingly, it appears that Golaghat district serves as the linking district where both the seeded and seedless fruits are present connecting the regions of Central Assam, North Assam, Lower Assam and Barak valley (Figure 10).

**Figure 9:**
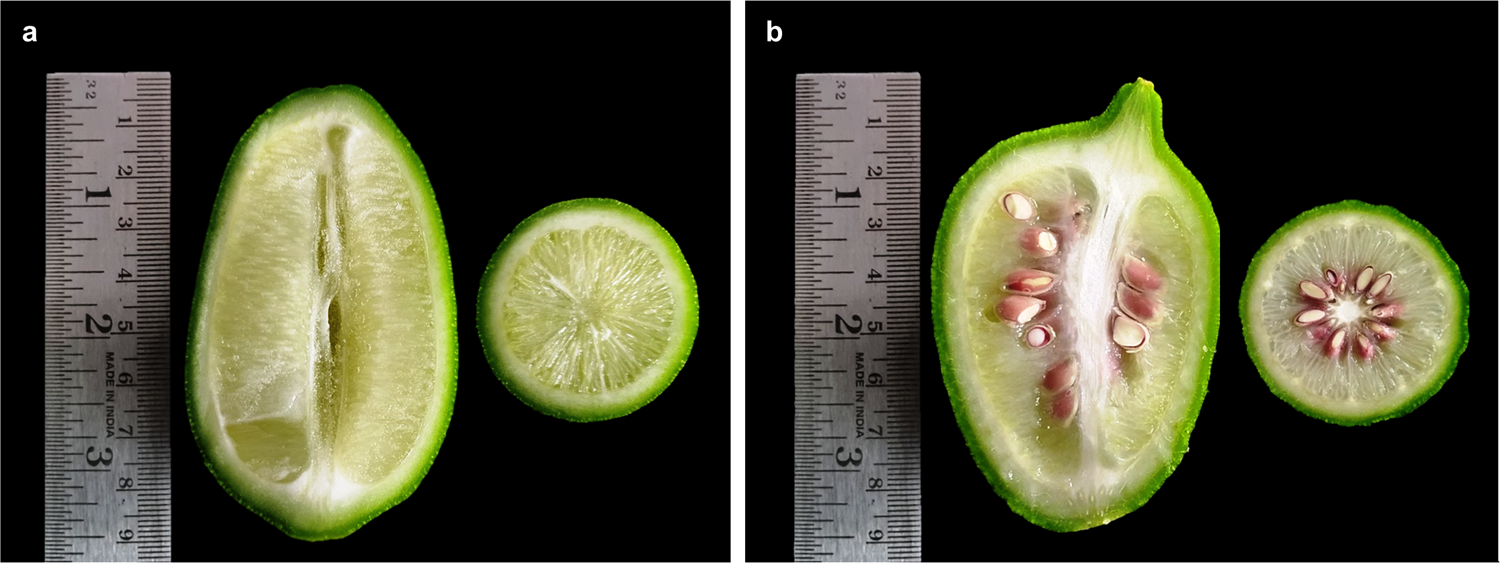
Assam lemon fruit; **a:** Seedless fruit, **b:** Seeded fruit

**Figure 10:**
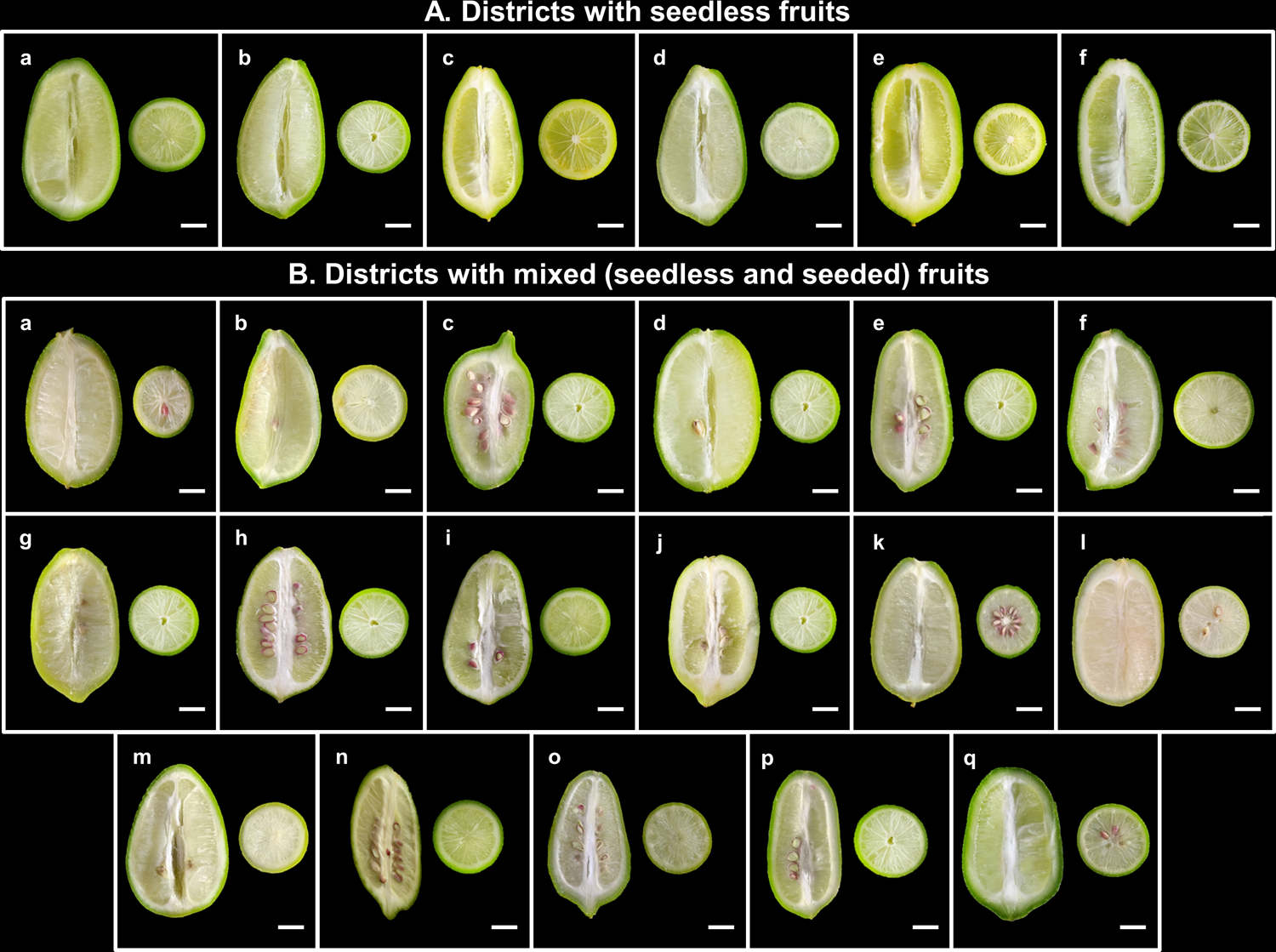
Variation in seed availability in Assam lemon fruits across different districts of Assam; **A:** Districts with seedless fruit (**a:** Control, **b:** Tinsukia, **c:** Dhemaji, **d:** Lakhimpur, **e:** Dibrugarh, **f:** Jorhat), **B:** Districts with mixed (Seedless and Seeded) fruits (**a:** Golaghat, **b:** Sonitpur, **c:** Karbi Anglong, **d:** Nagaon, **e:** Morigaon, **f:** Udalguri, **g:** Baksha, **h:** Nalbari, **i:** Barpeta, **j:** Kamrup Metro, **k:** Kamrup Rural, **l:** Bongaigaon, **m:** Kokrajhar, **n:** Dhubri, **o:** Dima Hasao, **p:** Cachar, **q:** Karimganj), Scale Bar: 1.5 cm

Further, during our comprehensive investigation of all 690 accessions (660 accessions collected from 132 populations of 22 districts, along with 30 accessions of one control population), we conducted a detailed analysis of the seeding character. Surprisingly, we discovered significant variation in the presence and quantity of seeds among the different districts of Central Assam, North Assam, Lower Assam and Barak Valley. Each district exhibited its own distinct seed count in the fruits where seeds were observed. While studying the seed count of Assam lemon fruits, we made interesting observations regarding different districts. Accessions from Golaghat, Sonitpur, Nagaon, Morigaon, and Nalbari districts consistently exhibited a similar pattern, with seed counts ranging from 0 to 5 seeds per district. On the other hand, when studying the accessions from Karbi Anglong, Udalguri, Baksha, Kamrup Metro, and Kamrup Rural districts, we found the seed counts ranging from 0 to 10 seeds per districts. Additionally, accessions from Barpeta, Bongaigaon, Kokrajhar, Dhubri, Dima Hasao, Cachar, and Karimganj districts also showed mixed seed counts, varying from 0 to >10 seeds per accession, however, certain accessions from the Dhubri and Cachar regions exhibited a notably high seed count, surpassing 20 seeds per accession (Table SF 4).

### PCoA Analysis

A Principal Coordinates Analysis (PCoA) was conducted to visualize the spatial patterns of morphological variation among populations and districts of Assam lemon. The results of the analysis showed that the populations from districts of Dhemaji, Tinsukia, Dibrugarh, Lakhimpur, Jorhat, and Dima Hasao shared highest morphological similarities with the control population (Figure 11). On the other hand, the populations from the districts of, Kamrup Metro, Kamrup Rural, Nalbari, Barpeta, Sonitpur, Nagaon, Morigaon, Baksha, Udalguri, Bongaigaon, Dhubri, Kokrajhar, Karbi Anglong, and Cachar exhibit greater dissimilarity from the control population in terms of morphological characteristics. Interestingly, the Ponkagaon (PGG) population of Golaghat district exhibited significant dissimilarity from the control population, while other Golaghat district populations maintained close similarities.

**Figure 11:**
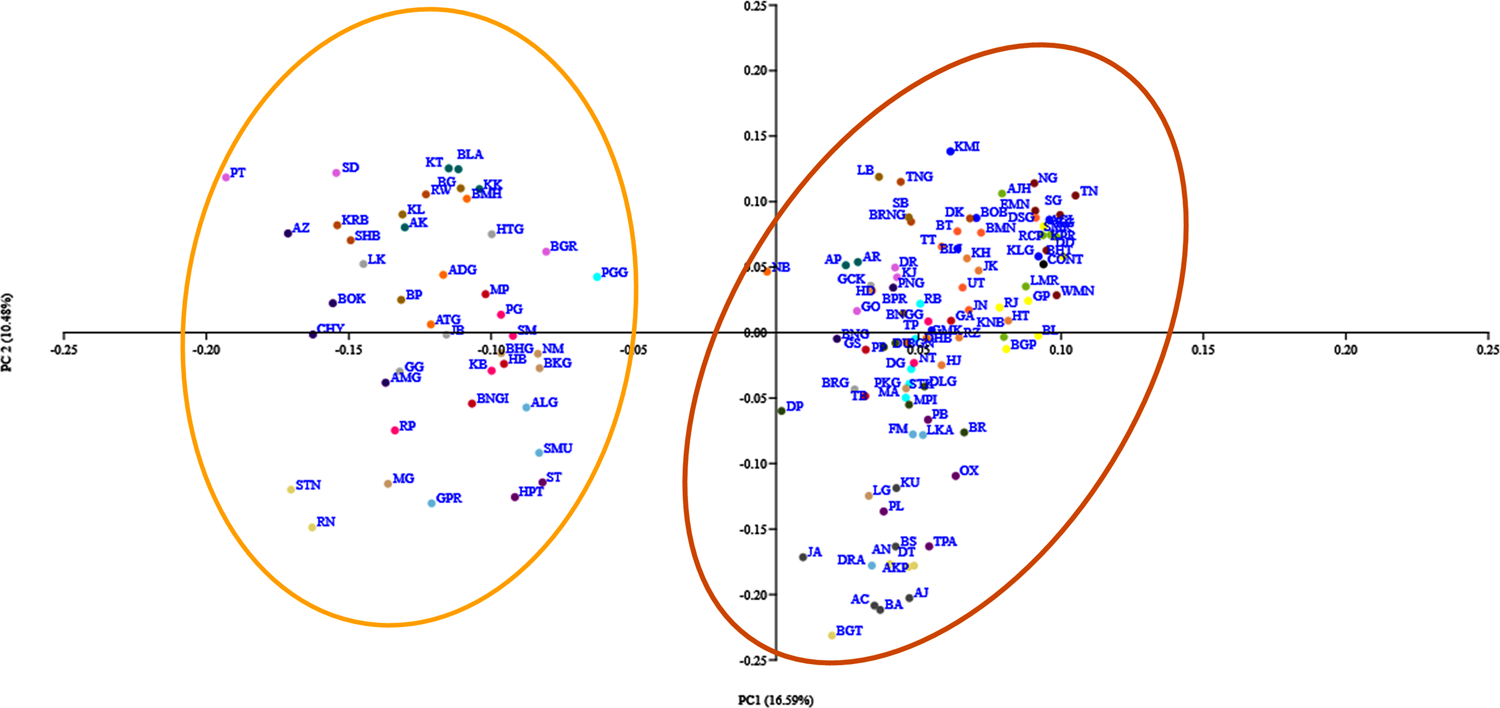
Principal Coordinate Analysis of Assam lemon using morphological characters

### Cluster Analysis

The dendrogram-based UPGMA algorithm analysis of morphological and seeding pattern data at the district level, led to identification of revealed two major clusters in the resulting dendrogram. Tn the dendrogram based on morphological data, it was obtained that the populations from Tinsukia and Dhemaji exhibited the close clustering with the control population (Figure 12).

**Figure 12:**
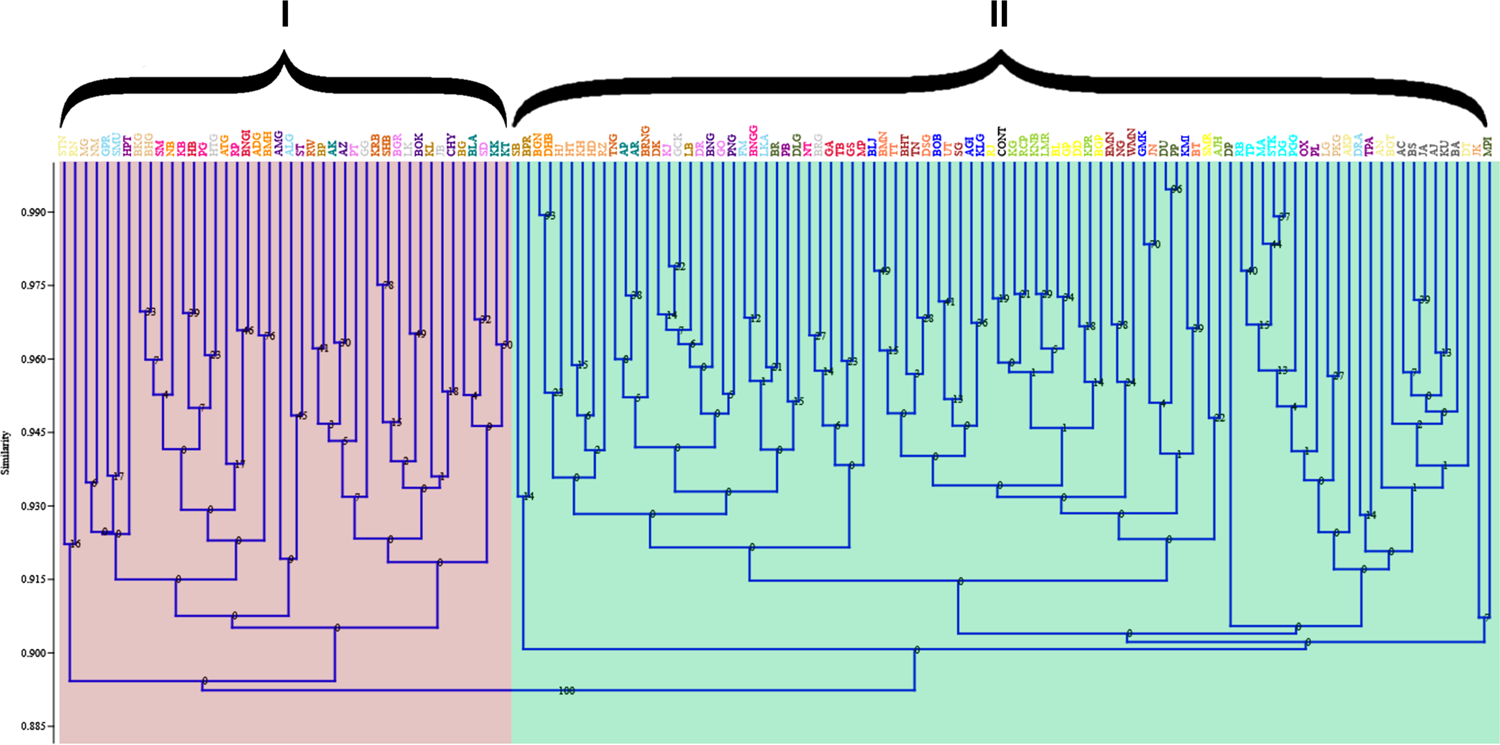
UPGMA analysis of Assam lemon populations using morphological characters. The alphabets bottom represents different populations code and their representative districts selected during the current study; **Control** (CONT), **Tinsukia** (BL, BGP, DD, GP, RJ, SMR), **Dhemaji** (KNB, LMR, KG, KPR, AJH, RCP), **Lakhimpur** (BLJ, BOB, GMK, AHI, KLG, KMI), **Dibrugarh** (EMN, NG, BHT, SG, TN, WMN), **Jorhat** (BMN, TT, JN, UT, DSG, BT), **Golaghat** (RB, PGG, MA, TP, STK, DG), **Sonitpur** (MG, LG, BKG, PKG, NM, BHG), **Karbi Anglong** (NB, ATG, ADG, BMH, BGN, DHB), **Nagaon** (RP, KB, BNGG, NT, PG, SM), **Morigaon** (BNGI, TB, HB, MP, GS, GA), **Udalguri** (RW, TNG, BRNG, KRB, SHB, DK), **Baksha** (BP, KL, SB, BG, BPR, LB), **Nalbari** (AK, BLA, KK, KT, AP, AR), **Barpeta** (PT, SD, DR, BGR, GO, KJ), **Kamrup Metro** (GG, JB, LK, BRG, HTG, GCK), **Kamrup Rural** (AZ, CHY, BOK, AMG, PNG, BNG), **Bongaigaon** (DU, PP, MPI, DP, DLG, BR), **Kokrajhar** (PL, TPA, HPT, ST, OX, PB), **Dhubri** (GPR, FM, DRA, ALG, LKA, SMU), **Dima Hasao** (HT, HJ, HD, KH, JK, RZ), **Cachar** (AN, BGT, STN, DT, RN, AKP), **Karimganj** (AC, AJ, BA, BS, JA, KU)

### Biochemical Analysis

The analysis of biochemical parameters in Assam lemon samples collected from various districts of Assam revealed substantial variation across the studied regions. Interestingly, we also observed a striking similarity between the biochemical parameters of the control populations and those from the districts of Dhemaji, Tinsukia, Lakhimpur, Dibrugarh, and Jorhat.

### pH

The results of the pH analysis of different districts in Assam along with the control population revealed that samples from the Cachar district had the highest pH (2.78 ± 0.01), while the samples from Lakhimpur district showed the lowest (2.27 ± 0.02) (Figure 13, Table SF 5).

**Figure 13:**
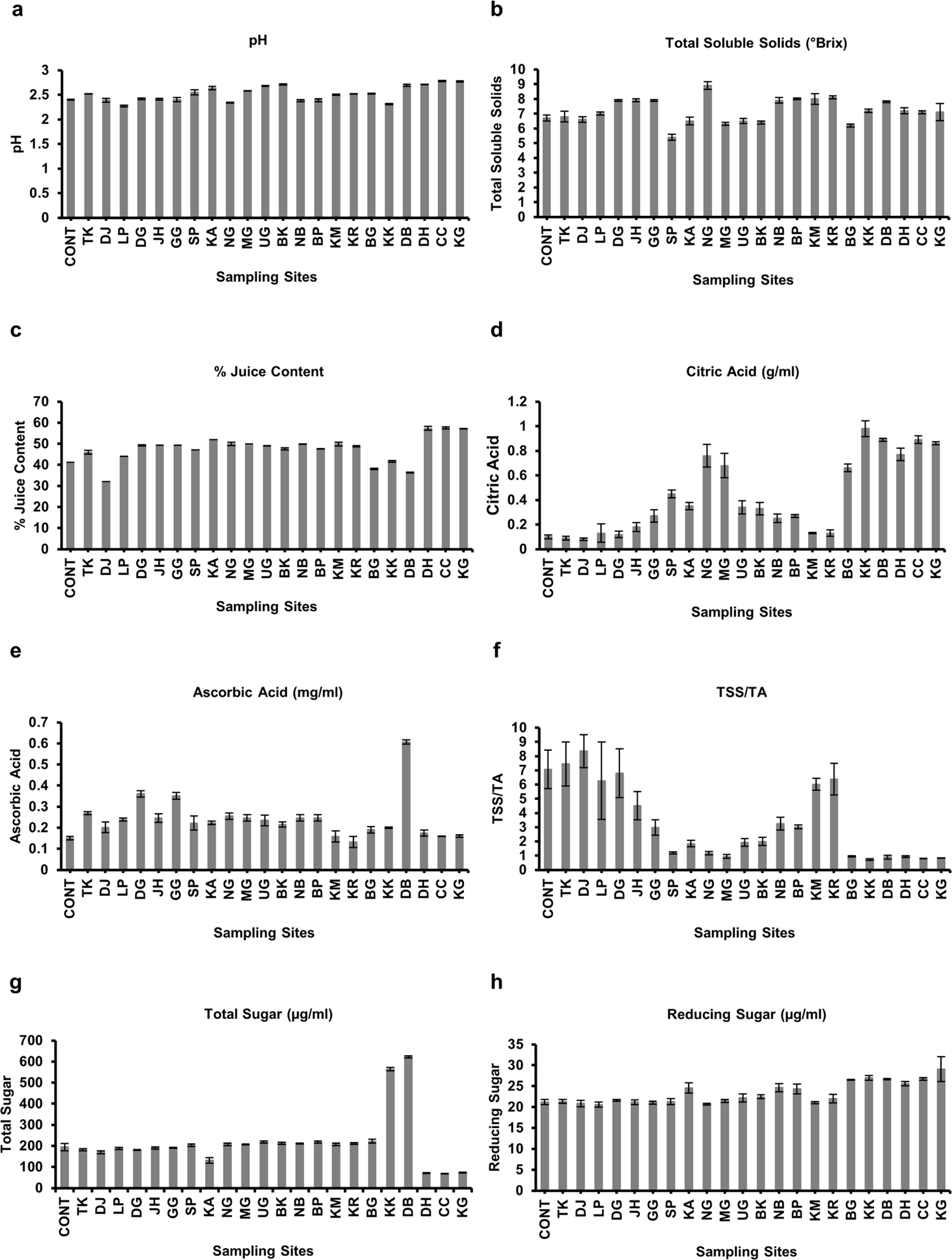
Biochemical characteristic of Assam lemon fruit juice across different districts of Assam; **a:** pH, **b:** TSS, **c:** % Juice Content, **d:** Citric Acid, **e:** Ascorbic Acid, **f:** TSS/TA, **g:** Total Sugar, **h:** Reducing Sugar

### % Juice Content

The % juice content analysis of different districts in Assam and the control population showed significant variations in the juice content of the samples. The samples from Cachar district had the highest % juice content (57.61 ± 0.523%), while, the samples from Dhemaji district exhibited the lowest % juice content (32.13 ± 0.036%) (Figure 13, Table SF 5)

### Total Soluble Solids

The results of the total soluble solids content analysis for different districts in Assam along with the control population revealed that the samples from Nagaon district had the highest total soluble solids (8.9 ± 0.27 °Brix), while Sonitpur had the lowest (5.4 ± 0.2 °Brix) (Figure 13, Table SF 5).

### Citric Acid

The results of the citric acid content analysis for different districts in Assam along with the control population revealed that Kokrajhar had the highest citric acid content (0.098 ± 0.0006 g/ml), while Dhemaji had the lowest (0.008 ± 0.0001 g/ml) (Figure 13, Table SF 5).

### Ascorbic Acid

The results of the ascorbic acid content analysis for different districts in Assam along with the control population revealed that Dhubri had the highest ascorbic acid content (0.607 ± 0.011 mg/ml), while Kamrup Rural had the lowest (0.132 ± 0.026 mg/ml) (Figure 13, Table SF 5).

### Total Sugar and Reducing Sugar

The results of the total sugar content esitimation for different districts in Assam, as well as the control population, revealed that Dhubri had the highest concentration of total sugar content (622.38 ± 5.646 µg/ml) and Cachar had the lowest (67.67 ± 0.97 µg/ml). Furthermore, results of the reducing sugar content analysis showed that Kokrajhar had the highest concentration of reducing sugar content (26.99 ± 0.6 µg/ml), while the lowest concentration was found in Lakhimpur (20.56 ± 0.6 µg/ml) (Figure 13, Table SF 5).

### Carotenoid Content

The results of the carotenoid estimation for the various districts in Assam along with the control population showed that the higest concentration of carotenoid on the peel of Assam lemon fruit was observed in the Nalbari district (2.64 ± 0.001 mg/g), while the lowest concentration was observed in Cachar district (1.17 ± 0.0058 mg/g) (Figure 14, Table SF 5).

**Figure 14:**
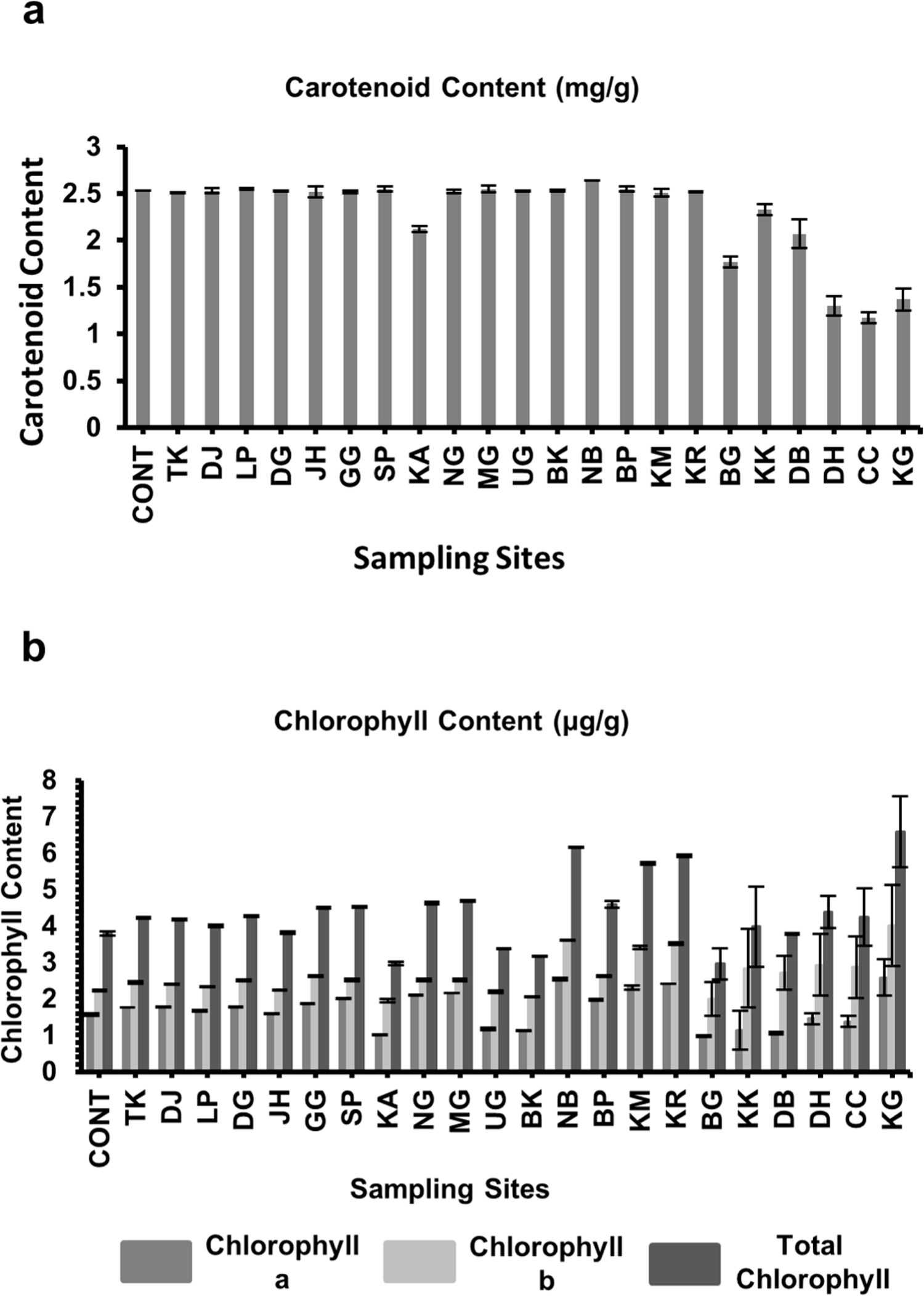
Biochemical characteristic of Assam lemon fruit juice across different districts of Assam; **a:** Carotenoid Content, **b:** Chlorophyll Content

### Chlorophyll Content

The results of the chlorophyll content estimation for the various districts in Assam, as well as the control population showed that the highest concentration of Chlorophyll a, Chlorophyll b, and the total chlorophyll content on the peel of Assam lemon fruit was observed in Karimganj district (2.58 ± 0.0496 µg/g, 4.01 ± 0.012 µg/g, 6.59 ± 0.0958 µg/g), and the lowest concentration was observed in Bongaigaon district (0.97 ± 0.0027 µg/g, 1.99 ± 0.046 µg/g, 2.96 ± 0.0434 µg/g) respectively (Figure 14, Table SF 5).

### Pectin Content

The results of pectin estimation for the various districts in Assam along with the control population revealed that Nalbari district had the highest concentration of pectin content on the peel of Assam lemon fruit (4.05 ± 0.282%), while Bongaigaon had the lowest concentration (2.14 ± 0.02%). On the other hand, Bongaigaon had the highest concentration of pectin content in the pulp of Assam lemon fruit (6.32 ± 0.02%), while Udalguri had the lowest (4.44 ± 0.572%) (Figure 15, Table SF 5).

**Figure 15:**
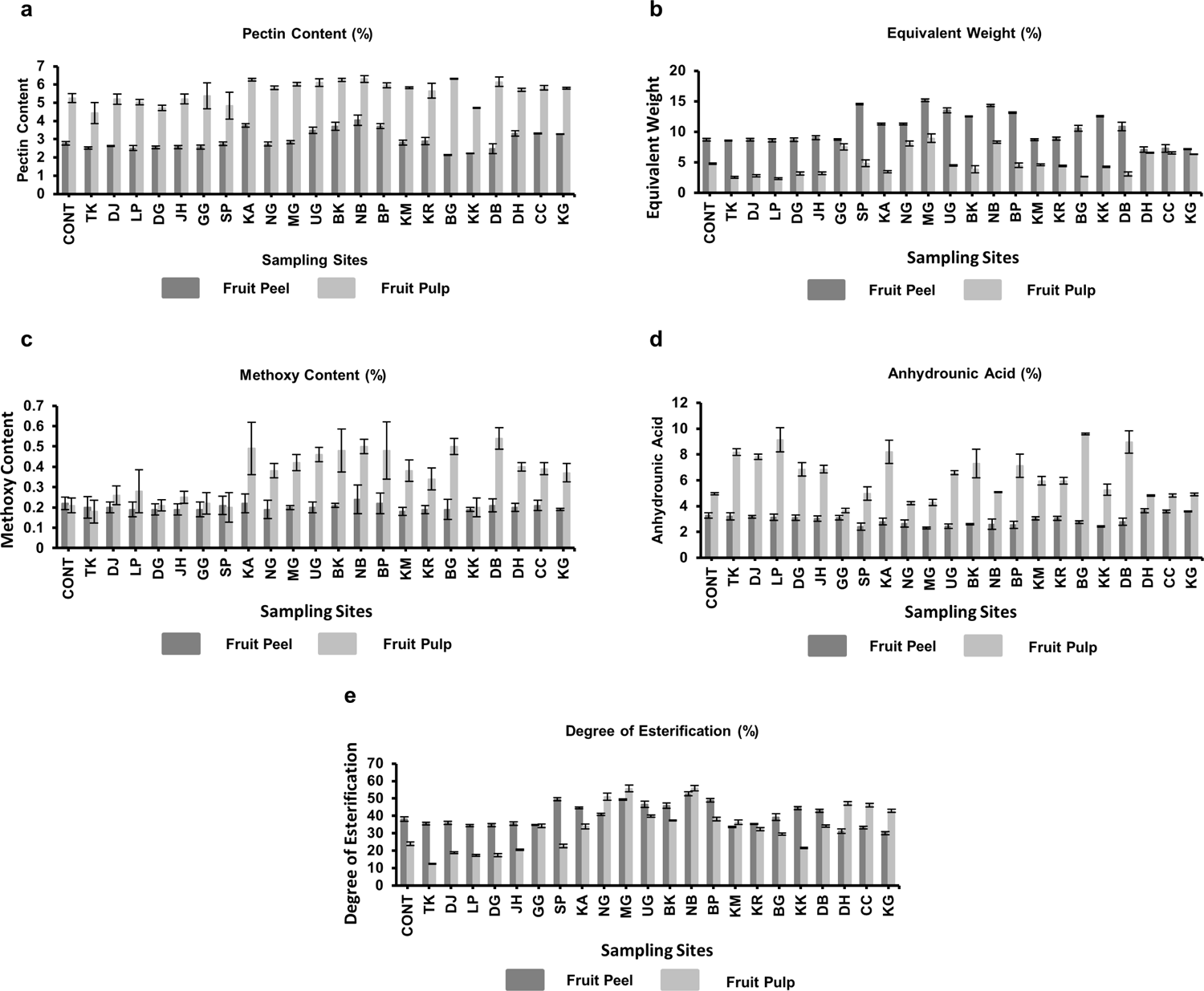
Chemical characterization of pectin obtained from Assam lemon fruit across different districts of Assam; **a:** Pectin Content, **b:** Equivalent Weight, **c:** Methoxy Content, **d:** Anhydrounic Acid, **e:** Degree of Esterification

### Equivalent Weight

The result of equivalent weight estimation for the various districts in Assam along with the control population showed that Sonitpur had the highest concentration of equivalent weight on the peel of Assam lemon fruit (15.17 ± 0.25%), while Dima Hasao had the lowest concentration (7.08 ± 0.447%). On the other hand, Morigaon had the highest concentration of equivalent weight in the pulp of Assam lemon fruit (8.96 ± 0.7%), while Lakhimpur had the lowest (2.31 ± 0.129%) (Figure 15, Table SF 5).

### Methoxy Content (MeO)

The result of methoxy content estimation for the various districts in Assam along with the control population showed that Nalbari had the highest concentration of methoxy content on the peel of Assam lemon fruit (0.24 ± 0.027%), while Morigaon had the lowest concentration (0.18 ± 0.02%). On the other hand, Dhubri had the highest concentration of methoxy content in the pulp of Assam lemon fruit (0.54 ± 0.5%), while Tinsukia had the lowest (0.18 ± 0.06%) (Figure 15, Table SF 5).

### Anhydrounic Acid (AUA)

The result of anhydrounic acid content estimation for the various districts in Assam along with the control population showed that Dima Hasao had the highest concentration of anhydrounic acid content on the peel of Assam lemon fruit (3.64 ± 0.15%), while Morigaon had the lowest concentration (2.3 ± 0.07%). On the other hand, Bongaigaon had the highest concentration of anhydrounic acid content in the pulp of Assam lemon fruit (9.59 ± 0.08%), while Tinsukia had the lowest (3.64 ± 0.17%) (Figure 15, Table SF 5).

### Degree of Esterification (DE)

The results of degree of esterification for the 22 selected different districts of Assam along with control population revealed that only Nalbari (52.61± 1.172%) district exhibited a high degree of esterification (>50%) in peel samples. In pulp samples, Nalbari, Morigaon, and Nagaon districts displayed the highest esterification levels at 55.95 ± 1.528%, 55.67 ± 1.924%, and 51 ± 1.978%, respectively. These findings emphasize the notable esterification concentrations in these districts’ peel and pulp samples compared to others, indicating potential variations in the composition of ester bonds in these regions (Figure 15, Table SF 5).

### Analysis of Variance (ANOVA)

During the investigation of variation of Assam lemon of different populations of Assam based on morphological and biochemical characters, we observed that there is a significant variation among the population of selected districts (p < 0.05), indicating a variation in the different morphological characters including seeding pattern and biochemical traits.

### Soil Nutrient Estimation

The soil nutrient assessment conducted for the current study revealed that the sampled populations exhibited a range from acidic to slightly alkaline. Most of the studied districts showed predominantly acidic soil, while populations from certain districts such as Lakhimpur, Golaghat, Sonitpur, Nagaon, Kamrup Metro, Kamrup Rural, Kokrajhar, and Dhubri displayed both acidic and alkaline pH levels. Interestingly, the control population and districts like Karbi Anglong, Dima Hasao, Cachar, and Karimganj showcased an alkaline soil nature.

The study also showed that the soil collected from all selected populations exhibited a significant presence of vital macro nutrients. The findings demonstrated that the soil in all populations of selected district contained a substantial amount (ranging from 0.505% to 1.5%) of Organic Carbon, except for the sites in Karbi Anglong, where the Organic Carbon concentration was comparatively lower (0.100% to 0.500%) compared to the populations of other districts. Moreover, the soil samples from the populations of Nagaon, and Dhubri along with the Control population, displayed a mixed concentration of Organic Carbon, ranging from 0.100% to 1.500%. Furthermore, current study also revealed that the soil samples from all populations in the selected districts exhibit a notable concentration of phosphate, ranging from 22 kg/ha to < 73 kg/ha, except for populations in Karbi Anglong district, which had phosphate levels > 22 kg/ha. In addition, the study findings revealed a significant presence of Potassium in the soil across most of the populations of selected districts, with concentrations ranging from 112 kg/ha to 392 kg/ha, except for the populations in Tinsukia, Dibrugarh, Dhemaji, Jorhat, Golaghat, Sonitpur, Morigaon, Kamrup Metro, and Kokrajhar. In these districts, the Potassium concentration (> 112 kg/ha) was relatively lower. On the other hand, the soil samples from Lakhimpur, Nagaon, Kamrup Rural, Nalbari, Bongaigaon, Dima Hasao, and Cachar displayed varying concentrations from low to high level of Potassium in the soil, ranging from > 112 kg/ha to 392 kg/ha. Additionally, the results show that, all the populations in the selected districts, including the Control population, displayed lower concentrations of Ammoniacal Nitrogen (15 kg/ha) and Nitrate Nitrogen (4 kg/ha to 10 kg/ha) in the soil (Table SF 6).

The study also revealed that in addition to macronutrients, the soil in different regions in the Assam is also abundant in various micronutrients (Copper, Zinc, Boron, Manganese, Iron, and Molybdenum) which plays a crucial role in the healthy growth and development of plants, even though they are required in smaller quantities compared to macronutrients. The findings demonstrated that the soil from all the selected districts contains a significant amount of Copper (ranging from 0.5 ppm to > 2 ppm) except some populations of Cachar and Dhubri districts along with all the populations of Karimganj district, where Copper content in the soil is relatively lower (ranging from 0 ppm to 0.5 ppm). It has also been noticed that the levels of Zinc in the soil samples are generally low, ranging from 0 to 0.5 ppm, except for specific areas in Tinsukia, Lakhimpur, and Cachar where the concentration of zinc was higher, ranging from 0.5 to 2 ppm. Again, Boron concentration in the soil samples exhibited significant variations across the sampled districts. Notably, Sonitpur, Nagaon, and Karbi Anglong districts displayed a significant presence (> 2 ppm) of Boron, whereas soil samples from the Cachar district and the Control population showed considerably lower Boron levels (ranging from 1 ppm to 2 ppm). In contrast, the soil samples of the remaining districts exhibited a diverse range of Boron concentrations, varying from 0.1 ppm to > 2 ppm. A notable variation in Manganese concentration was observed among all the populations of selected districts, demonstrating high Manganese levels (> 4 ppm) in the accessions of Tinsukia, Dhemaji, Lakhimpur, Udalguri, Baksha, Nalbari, and Barpeta, while Sonitpur, Nagaon, Kokrajhar, and Karbi Anglong exhibited comparatively lower concentrations (ranging from 2 ppm to 4 ppm) in contrast to the aforementioned districts in Assam. The concentration of Manganese were however found to be significantly low (0 ppm to 2 ppm) in the populations of Dibrugarh, Jorhat, Golaghat, Morigaon, Kamrup Metro, Kamrup Rural, Bongaigaon, and the Control population. Further, the soil samples from the populations of Dima Hasao, Cachar, Karimganj, and Dhubri displayed a diverse range of Manganese concentration, ranging from 0.2 ppm to > 4 ppm. Additionally, the soil samples from Dhemaji, Lakhimpur, Dibrugarh, Jorhat, Golaghat, Sonitpur, Udalguri, Baksha, Nalbari, Barpeta, Karbi Anglong, Nagaon, Morigaon, Kamrup Metro, Kamrup Rural, and Karimganj along with the control population displayed a significant concentration (ranging from 3 ppm to > 6 ppm) of Iron in the soil, whereas samples from all the populations in Dhubri were found to be significantly low in Iron concentration (ranging from 1 ppm to 3 ppm). Further, the soil samples collected from Tinsukia, Dima Hasao, Cachar, and Kokrajhar exhibited a wide range of Iron concentrations, ranging from 1 ppm to > 6 ppm. The concentration of Molybdenum however remained consistently low (ranging from 0 ppm to 1 ppm) across all districts, including in the control population (Table SF 6).

## Discussion

Morphological characters analysis plays a crucial role in the investigation of plants, enabling researchers to explore the vast array of variations in a plant species (Perez-Nicolas et al. 2020). By closely examining the morphological characteristics of plants, valuable insights can be gained into the diverse traits and variations within the different plant species, cultivars, and varieties but also within the same cultivar and variety across different regions (Qi et al. 2020; Alcantara-Ayala et al. 2020; Cabrera-Toledo et al. 2020). This comprehensive analysis helps researchers better understand the intricate details of plant morphology and facilitates advancements in plant breeding, taxonomy, and overall plant science (Alcantara-Ayala et al. 2020). The morphological variation observed among Assam lemon populations from different districts of Assam provides valuable insights into the morphological variation within the cultivar. During the investigation of variation of the morphological characters of the different populations, we observed that there is a significant variation in several morphological characters including fruit weight, fruit length, fruit diameter, pulp to peel ratio and others, among the 132 populations of the 22 districts along with the Control populations, thus indicating variation in important fruit characters of Assam lemon fruits. Specifically, the flowering pattern of Assam lemon collected from Upper Assam districts (except Golaghat) showed similarity with the findings of Bhattacharya and Dutta (1956), however, the concentration of unisexual flowers increased from Golaghat district to Central, North, and Lower Assam, and Barak Valley and was found to be of almost equal concentration to bisexual flowers.This variation could be attributed to various factors, including environmental conditions, soil composition, and farmer practices specific to each region (Liliane and Charles 2019). The ANOVA analysis revealed a statistically significant difference (p < 0.05) among the populations, indicating a notable variation in morphological characters (Andrade et al. 2019). This finding highlights the presence of distinct morphological variations within the Assam lemon cultivar across different populations, emphasizing the significance of studying and understanding these variations in the context of plant morphology and genetic diversity. Similar investigation was also observed on *C. nobilis* Lour (Chan et al. 2022), *Cynodon dactylon* (L.) Pers. (Wang et al. 2020), *C. reticulate* Blanco (Dorji and Yapwattanaphun 2011) which highlighted the differences in morphological traits across different populations. These findings have practical implications for growers and breeders as they can select suitable populations with desired morphological characters for commercial cultivation and breeding programs.

Also, the PCoA analysis of the population utilizing morphological characters provided compelling evidence of high morphological divergence among the Assam lemon populations. The results indicated the presence of two distinct and well-defined groups that vary from each other in terms of their morphological traits. This emphasizes the significance of considering the morphological diversity within the Assam lemon cultivar, as it holds implications for genetic differentiation, adaptation, and potential breeding strategies (Wang et al. 2015). These findings also open up avenues for further investigation into the underlying factors (such as environmental influences, genetic factors, or geographical factors) contributing to the observed morphological variations, thus can enhance our understanding of the complex dynamics within the Assam lemon populations (Pacheco-Hernandez et al. 2021). Similar observations were also documented for UPGMA clustering analysis, where a total of 133 populations formed two distinct clusters. Notably, the second cluster displayed further subdivision into two sub-clusters. Additionally, the UPGMA clustering analysis highlighted that the populations from Dhemaji and Tinsukia districts exhibited the highest degree of morphological similarity with the control population. Interestingly, in a recent study conducted by Ahmed et al. in 2023, they also discovered similar outcomes while investigating the genetic diversity of Assam lemon using ISSR marker system where populations from Dhemaji and Tinsukia districts showed close resemblance with the control population as obtained from UPGMA clustering analysis. This observation could be attributed to relatively fewer mutations occurring over time in these populations compared to other populations studied. In contrast, populations from Kokrajhar, Bongaigaon, Dhubri, Cachar, and Karimganj exhibited significant morphological differentiation from the Control population. Similar observations has been made in few other species such as *Cyclamen sp*. (Cornea-Cipcigan et al. 2023), *Cichorium sp*. (El-Taher et al. 2023), *C. Maxima* (Burm.) Merr. (Susandarini et al. 2013). In 2023, genetic diversity investigation on Assam lemon populations conducted by Ahmed et al. revealed that the aforementioned districts were genetically closer to the Control population compared to populations from other districts (Ahmed et al. 2023). This implies that genetic variations or mutations could be one of the reasons for the observed variation in the presence or absence of seeds in different regions of Assam (Gnan et al. 2014).

Furthermore, during the investigation of seed count among different populations in selected districts, we have observed significant variation in the number of seeds. Specifically, the populations of Dhemaji, Tinsukia, Dibrugarh, Lakhimpur, and Jorhat, along with the Control population, exhibited a seedless nature in their fruits which is its characteristic nature for which this cultivar was introduced and became popular (Bhattacharya and Dutta 1956). However, the populations starting from Golaghat districts towards Central Assam, North Assam, Lower Assam and Barak Valley districts showed a mixed response, with some accessions being having seeded and others seedless fruits. Interestingly, the results of the seeding pattern resemble with the variation in flowering nature that we observed during the investigation of morphological variation. This might suggests that the flowering nature variation along with the nature of cultivation practices might be a contributing factor to the differences in seed development in Assam lemon, also suggesting that Assam lemon might be a self-incompatible cultivar having tendency towards natural hybridization (Khan et al. 2004, Velasco and Licciardello 2014, Wu et al. 2014).

Additionally, the number of seeds found in Assam lemon fruits across different accessions was studied. The study observed variations in seed counts among different accessions, with specific patterns emerging in different districts with variations ranging from no seeds to more than 10 seeds. Seed counts ranged from 0 to 5 in certain districts (Golaghat, Sonitpur, Nagaon, Morigaon, and Nalbari), 0 to 10 in others (Karbi Anglong, Udalguri, Baksha, Kamrup Metro, and Kamrup Rural), and more than 10 in yet others (Barpeta, Bongaigaon, Kokrajhar, Dhubri, Dima Hasao, Cachar, and Karimganj). The variations in seed counts among different accessions of Assam lemon fruits, coupled with specific patterns observed in different districts, indeed highlight the complexity of factors influencing seed patterns. The combination of environmental conditions, genetic mutations, and soil nutrient variations across districts may contribute to the observed diversity in seed counts. It was previously reported that the environmental factors and genetic mutations act as a potential influencer on the seeding pattern of fruits (Cao et al. 2020). Also, the variation in the soil nutrients in different districts of Assam may be a reason, which contributes to the differences in seeding patterns among the accessions in the current study (Gnan et al. 2014). Furthermore, the inherent propensity of *Citrus* species, including Assam lemon, for natural hybridization could potentially account for variations in seed formation and seed counts among different populations over time (Neri et al. 2018). Given the contemporary emphasis on seedless fruit in response to consumer preferences, modern farmers and breeders are actively engaged in the development of diverse seedless fruit varieties (Premachandran et al. 2019). The investigation into the variations in the seeding pattern of Assam lemon holds the potential to significantly impact cultivation techniques, aligning them with consumer desires for enhanced fruit quality (Janeczko and Timmons 2019). By catering to demands for high yield, substantial juice content, and the absence of seeds, the outcomes of this study can play a pivotal role in guiding cultivation practices and meeting the evolving expectations of consumers (Bhandari et al. 2022).

Apart from the above mentioned observations, variation in different biochemical parameters was found in the fruits of Assam lemon across the selected districts. A significant variation in pH concentration was observed in the samples from Karbi Anglong, Sonitpur, Baksha, Udalguri, Morigaon, Dhubri, and the Barak Valley districts (Dima Hasao, Cachar and Karimganj) where the pH was shown to be higher (>2.5) as compared to other districts and the Control population. The change in pH may be due to the variation in agro-environmental factors, such as soil nutrient composition, rainfall, and temperature, which can influence pH levels in fruits (Etienne et al. 2013). The current investigation also revealed that the samples from Nagaon, Kamrup Metro, and Kamrup Rural districts e TSS ≥, w important biochemical parameter used in various processing applications, such as juicing, canning, and making jams, jellies, and others (Electronmachine 2017). The high TSS concentrations indicate an increased content of dissolved sugars, organic acids, and other soluble compounds in Assam lemon samples of Nagaon, Kamrup Metro, and Kamrup Rural districts (Li et al. 2021). Environmental conditions, including sunlight, rainfall, temperature, soil nutrients, and agricultural practices have been reported to affect TSS values in fruits (Porter et al. 1940). These factors could be the reason for variation in TSS concentrations observed across different regions of Assam (Porter et al. 1940).

Assam lemon is also rich in juice content as compared to other different lemon varieties due to its large size (Barua and Bharadwaj 2017). It was also revealed that there is a significant variation in the % of juice content among different regions of Assam. All the districts along with the control population exhibited % juice content below 50%, except for Morigaon, Karbi Anglong, and the districts of the Barak Valley, which displayed a juice content exceeding 50%. Interestingly, the results also revealed that, although the populations from Dhemaji and Tinsukia districts exhibited resemblances with the control population in terms of morphological and other biochemical attributes, a notable difference emerged in terms of % juice content. Specifically, the population from the Dhemaji district displayed a lower % juice content (<40%) at 32.13 ± 0.036% in comparison to both the control population and the Tinsukia district population (>40%) at 41.24 ± 0.071% and 45.92 ± 0.92% respectively. The observed variations in juice content across different regions can be attributed to a range of regional factors, such as climate, soil composition, agricultural practices, and even genetic variations (Garcia-Munoz et al. 2021; Tasisa et al. 2018). Further, during the current investigation, an interesting observation regarding the concentration of citric acid in Assam lemon across different districts was noted, it was observed that the districts of Dhemaji, Tinsukia, and Dibrugarh in Upper Assam and the control population had the lowest concentration of citric acid. However, as we moved towards Jorhat and Golaghat, the concentration of citric acid gradually increased. The districts in the Barak Valley exhibited the maximum concentration of citric acid. This variation in citric acid content suggests differences in the enzymatic activity responsible for citric acid synthesis during fruit development (Mukhim et al. 2015). In addition to citric acid, we also examined the ascorbic acid content in the Assam lemon samples. Surprisingly, it was found that all the districts, along with the control population, had ascorbic acid content below 0.4 mg/ml, except for the samples from the Dhubri district, which displayed an ascorbic acid content of 0.607 mg/ml. The variation in acid content among the Assam lemon fruits could be attributed to environmental conditions, including temperature, sunlight exposure, rainfall, and other factors (Carr and Rowe 2020). It has been reported that the ascorbic acid content of *Citrus* fruits is not stable and can vary due to enzymatic loss, where L-ascorbic acid is converted to 2-3- deoxy-L-gluconic acid (Gurung et al. 2022). Additionally, the utilization of ascorbic acid in metabolic processes might contribute to its decrease during the maturation process (Padayatty and Levine 2016).

On the other hand, it was observed that the fruits obtained from Dhemaji, Tinsukia, Dibrugarh, Jorhat, and Golaghat, as well as Kamrup Metro and Kamrup Rural districts, exhibited high level of TSS/TA compared to samples from other districts. Fruits from these regions displayed higher levels of TSS and a significant decrease in acid accumulation, which could be the contributing factors to the decline in the TSS/TA ratio (Mukhim et al. 2015). Further, during the current investigation of Assam lemon fruit juice, a noteworthy finding emerged regarding the relationship between sugar concentration and pH, we observed a negative correlation between these two variables across most of the samples analyzed. This implies that as the sugar concentration increased, there was a corresponding decrease in the pH of the juice (De Montfort University 2019). This intriguing pattern might be due to the accumulation of organic acids within the juice (Liao et al. 2019). These organic acids not only impact the sugar concentration but also play a crucial role in determining the overall pH of the fruit juice (Huang et al. 2021). Hence, the observed variations in sugar concentration may be attributed to the influence of these organic acids, which subsequently affect both the sugar concentration and the pH levels of the juice (Huang et al. 2021; Liao et al. 2019). Additionally, while studying the carotenoid and chlorophyll content in samples of selected districts along with the samples from control population, a negative correlation between the carotenoid and chlorophyll content was observed. The environmental factors such as light intensity, temperature, and nutrient availability may be the causes which influenced the pigment composition of Assam lemon peel (Jing et al. 2020). Low chlorophyll levels and high carotenoid levels may be favored by specific environmental conditions that promote carotenoid synthesis and accumulation (Lado et al. 2019).

Pectin, a crucial ingredient used in various food products, including jams, jellies, and low-calorie foods, was found to be abundant in both the peel and pulp of the selected samples from different districts, including the control population (Kumar et al. 2018). However, it was noted that the pectin concentration was consistently high in the pulp compared to the peel. Interestingly, when comparing the pectin concentration among districts, it was found that samples from Upper Assam and the control population exhibited lower pectin concentrations compared to samples from other districts. Environmental conditions and genetic mutations might be the attributing factors to these variations, which have also been previously reported to influence pectin production (Du et al. 2020; Venkatanagaraju et al. 2019). Pectin can be categorized as either high-methoxy pectin or low-methoxy pectin based on its esterification extent (Khamsucharit et al. 2017). High-methoxy pectin is known for its sensitivity to acid conditions and typically requires a substantial amount of sugar to function as a thickening and gelling agent (Akin- Ajani and Okunlola 2021). On the other hand, low-methoxy pectin has gained prominence in the food industry, particularly for jam production, as it can form a gel with less reliance on sugar (Akin-Ajani and Okunlola 2021). To ensure the purity of the extracted pectin, we also evaluated the anhydrounic acid content in both the peel and pulp of Assam lemon samples (Khamsucharit et al. 2017). Notably, the pulp displayed the highest anhydrounic acid content, followed by the peel. Findings from the current study indicated that samples from the control population, Upper Assam, Barak Valley, Bogaigaon, Kokrajhar, and Dhubri districts have highest pectin purity compared to samples from other districts. The present investigation also delved into the esterification level of pectin, specifically focusing on the concentration of methylesters, which plays a significant role in determining the applicability of pectin in various food industries (Vanitha and Khan 2019). The present study also made an interesting observation regarding the pectin esterification levels. Notably, the peel samples from Nalbari and the pulp samples from Morigaon and Nalbari exhibited high concentrations of methylesters, indicating the presence of high-methoxy pectin. In contrast, the samples from all other districts, including the Control population, displayed low concentrations of methylesters, suggesting the prevalence of low-methoxy pectin. Factors such as climate, soil composition, and agricultural practices specific to each region may contribute to these variations (Etienne et al. 2013). These factors may collectively influence the physiological characteristics of Assam lemon trees, leading to variations in fruit development and juice production (Zhang et al. 2022; Wang et al. 2015). The ANOVA analysis revealed a statistically significant difference (p < 0.05) among the different districts, indicating a notable variation in biochemical attrubutes characters (Andrade et al. 2019).

The soil composition and nutrient availability play a pivotal role in shaping the morphological characters of plants, including various fruit traits such as length, diameter, peel thickness, seed number, and more (Schjoerring et al. 2019). During the current investigation, soil nutrient analysis from 133 populations revealed a significant variation not only in the nutrient content but also in the soil nature itself. This variation in soil characteristics is likely to be a crucial factor contributing to the observed variations in different morphological characters of Assam lemon within the selected populations (Bano et al. 2019). It’s important to note that soil fertility and nutrient content can vary widely based on geographical and environmental factors. Therefore, regular soil testing and analysis are essential for farmers to make informed decisions regarding nutrient management and crop production.

## Conclusion

The findings from the current investigation highlight notable differences in morphological, seeding pattern, and biochemical traits among Assam lemon accessions. The observed pattern of differentiation is unlikely to have occurred randomly within a relatively short span of 67 years, which is the time frame for clonal propagation from a single seedling, which showed similar results with the findings in Assam lemon by Ahmed et al in 2023 (Ahmed et al. 2023). These variations strongly indicate that the sampled accessions are not clonal propagules derived from a single parent plant. Further, the study suggests a plausible scenario where the sampled accessions have independently arisen in various locations as chance propagules and have subsequently been promoted for cultivation, both commercially and domestically. This hypothesis gains support from the fact that the control population exhibits closer alignment, in terms of seeding pattern, morphological and biochemical characteristics, with the districts from Upper Assam compared to the other selected districts in Assam. In light of these findings, it is crucial to consider the preservation of each extant population as an integral part of a comprehensive management strategy. Preserving these populations is essential to safeguard their unique qualities and characteristics, which may have significant implications for future breeding programs, conservation efforts, and the cultivation of Assam lemon for various purposes. Additionally, the current study will also assist the farmers in cultivation, and improved management practices of Assam lemon, ensuring its continued success in agricultural landscape of Assam. Furthermore, the knowledge generated from the current study will also serve as a foundation for future research and the development of evidence-based recommendations and guidelines for successful Assam lemon cultivation in Assam and beyond.

## Author contributions

Original draft preparation, investigation, acquisition of data, analysis, and interpretation of data was performed by Suraiya Akhtar, and Raja Ahmed. Suraiya Akhtar and Raja Ahmed contributed equally to this work and share first authorship. The article was reviewed by Ankur Das and Khaleda Begum. Assess to control samples for the experiment was facilitated by Dr. Sarat Saikia. The conception, study design, supervision, critical review and the article revision was done by Sofia Banu. All authors read and approved the final manuscript. The authors also declare no conflict of interest.

## Acknowledgement

The authors are indebted to Gauhati University, and Horticultural Research Station, Kahikuchi for providing the technical facility and Control samples for the current investigation. The authors acknowledge the help rendered by all individuals (specially Mayuri Das, Bishmita Boruah, Pulakeswar Bsumatary, Barbi Bhuyan, Hungma Boro, Swati Gupta, Bhaswati Ramacharya, Rajashri Sarma, Madhuchanda Sarma, Suman Goswami, Ahmed Sarif Laskar, Mafizur Rahman, Hafizur Rahman, Eunus Ali Mondal, Shohid Ali Mondal, Musadiqque Mahfuz Ahmed Mahfuz Ahmed, Rabi Ali, Swmkhwr Shaan Swargiary, Sajid Ali, Aziz Ali Mondal, Swapna Bharali, Ahmed Hussain, Akhil Kalita, Rulee Laskar, Subham Roy, Kalpajyoti Dutta, Sopon Neog, Dr. Fakaruddin Ali Ahmed, Ponkaj Thengal, Manjuma Begum, Hiranya Bhatta, Fatima Siddiqua, Aman Gupta, Abdul Ali, Asiquer Rahman, Rejakul Ahmed, Bidyut Das, Karabi Baruah, Pranjit Kalita, Malaya Ghosh, Moni Kalita, Ananya Dutta, Farnaz Irin Alom, Jutika Borah, Sonali Saha, Nilofer Ahmed, and Rahul Thaosen) who helped in the collection of different accessions across different districts of Assam. Also the authors thanked Dr. Mrinal Baruah for his help in preparation of the map. The authors also acknowledge Department of Biotechnology, Government of India for providing the financial aid vide sanction number BT/PR21144/BPA/118/203/2016 under Task Force scheme.

## Statement and Declaration

### Competing Interest

The authors have no relevant financial or non-financial interests to disclose.

### Data availability statement

Data is contained within the article and the supplementary files.

## Funding

This work was supported by Department of Biotechnology, Government of India under Task Force (sanction number BT/PR21144/BPA/118/203/2016).

## Supplementary File Legends

**Table SF 1:** Details of Assam lemon accessions collected from different districts of Assam

**Table SF 2:** Variation in morphological characters of Assam lemon accessions across different districts of Assam

**Table SF 3:** Variation in flowering characters of Assam lemon accessions across different districts of Assam

**Table SF 4:** Variation in fruit characters and seed characters of Assam lemon accessions across different districts of Assam

**Table SF 5:** Variation in biochemical traits of Assam lemon across different districts of Assam

**Table SF 6:** Estimation of micronutrients present in the soil of Assam lemon population across different districts of Assam

**Table SF 7:** Estimation of macronutrient present in the soil of Assam lemon population across different districts of Assam

## Notes

### Competing Interest Statement

The authors have declared no competing interest.

## References

Ahmed, R., Akhtar, S., Das, A., Begum, K., Kashyap, K., Banu, S., 2023. Delineating the degree of genetic divergence within Assam lemon (*Citrus limon*, I G R Evol. 10.1007/s10722-023-01606-8.

Akin-Ajani, O.D., Okunlola, A., 2021. Pharmaceutical applications of pectin, in: Masuelli, M.A. (Ed.), Pectins. IntechOpen., London, pp 1–17. 10.5772/intechopen.100152.

Alaida, M.F., Aldhebiani, A.Y., 2022. Comparative study of the morphological characteristics of *Phoenix dactylifera* L. cultivars in Al-Madinah Al-Munawarah-Saudi Arabia. BMC Plant Biol 22, 461. 10.1186/s12870-022-03841-0.

Alcantara-Ayala, O., Oyama, K., Rios-Munoz, C.A., Rivas, G., Ramirez-Barahona, S., Luna-Vega, I., 2020. Morphological variation of leaf traits in the *Ternstroemia lineata* species complex (Ericales: Penthaphylacaceae) in response to geographic and climatic variation. PeerJ 8, e8307. 10.7717/peerj.8307.

Andrade, C., 2019. The P value and statistical significance: misunderstandings, explanations, challenges, and alternatives. Indian J Psychol Med 41(3), 210–215. 10.4103/IJPSYM.IJPSYM_193_19.

Bano, C., Amist, N., Singh, N.B., 2019. Morphological and anatomical modifications of plants for environmental stresses, in: Roychoudhury, A., Tripathi, D. (Eds.), Molecular plant abiotic stress: Biology and biotechnology. John Wiley and Sons, Ltd., New Jersey, 29–44. 10.1002/9781119463665.ch2.

Bardel-Kahr, I., Schliep, K., 2023. Phylogenetic trees from morphological data. Accessed February 2023. https://cran.r-project.org/web/packages/phangorn/vignettes/Morphological.html.

Barua, B.C., Bharadwaj, S., 2017. Assam Lemon – A prospective npd initiative aimed at global market positioning. Int J Res 4(14), 727–734.

Bhandari, S.R., Rhee, J., Choi, C.S., Jo, J.S., Shin, Y.K., Song, J.W., Kim, S.H., Lee, J.G., 2022. Morphological and biochemical variation in carrot genetic resources grown under open field conditions: the selection of functional genotypes for a breeding program. Agron 12(3): 553. 10.3390/agronomy12030553.

Bhattacharya, S.C., Dutta, S., 1956. Classification of Citrus fruits of Assam. Scientific Monograph No. 20. I.C.A.R, New Delhi.

Brima, E.I., Abbas, A.M., 2014. Determination of citric acid in soft drink, juice drinks and energy drinks using titration. Int J Chem Stud 1(6), 30–34. 10.13140/2.1.1882.6886.

Buckan, D.S., 2015. Estimation of glycemic carbohydrates from commonly consumed foods using modified anthrone method. Indian J Appl Res 5(3), 45–47.

Budiarto, R., Poerwanto, R., Santosa, E., Efendi, D., 2021. Morphological evaluation and determination keys of 21 Citrus genotypes at seedling stage. Biodivers 22(3), 1570–1579. 10.13057/biodiv/d220364.

Cabrera-Toledo, D., Vargas-Ponce, O., Ascencio-Ramirez, S., Valadez-Sandoval, L.M., Perez-Alquicira, J., Morales-Saavedra, J., Huerta-Galvan, O.F., 2020. Morphological and genetic variation in monocultures, forestry systems and wild populations of *Agave maximiliana* of Western Mexico: implications for its conservation. Front Plant Sci 11, 817. 10.3389/fpls.2020.00817.

Cao, J., Chen, L., Wang, J., Xing, J., Lv, X., Maimaitijiang, T., Lan, H., 2020. Effects of genetic and environmental factors on variations of seed heteromorphism in *Suaeda aralocaspica*. AoB Plants 12(5), plaa044. 10.1093%2Faobpla%2Fplaa044.

Carr, A.C., Rowe, S., 2020. Factors affecting Vitamin C status and prevalence of deficiency: a global health perspective. Nutr 12(7), 1963. 10.3390/nu12071963.

Chan, S.R.O.S., Achmad, B.S., Ferdinant., 2022. Morphological characterization of Gunung Omeh Citrus (*Citrus nobilis* Lour) in Guguak District, Lima Puluh Kota Regency. IOP Conf Ser: Earth Environ Sci 1097, 012032. 10.1088/1755-1315/1097/1/012032.

Cornea-Cipcigan, M., Pamfil, D., Sisea, C.R., Margaoan, R., 2023. Characterization of *Cyclamen* genotypes using morphological descriptors and DNA molecular markers in a multivariate analysis. Front Plant Sci 14, 1100099. 10.3389/fpls.2023.1100099.

De Montfort University, 2019. Scientists find a way to reduce sugar in drinks. Accessed June 2023. https://www.dmu.ac.uk/research/research-news/2019/july/scientists-find-a-way-to-reduce-sugar-in-drinks.aspx#:~:text=%E2%80%9CSugar%20is%20being%20used%20to,acidic%20pH%20level%20of%202.5.%E2%80%9D.

Dorji, K., Yapwattanaphun, C., 2011. Assessment of morphological diversity for local mandarin (*Citrus reticulata* Blanco.) accessions in Bhutan. J Agric Technol 7(2), 485–495.

Du, J., Kirui, A., Huang, S., Wang, L., Barnes, W.J., Kiemle, S.N., Zheng, Y., Rui, Y., Ruan, M., Qi, S., Kim, S.H., Wang, T., Cosgrove, D.J., Anderson, C.T., Xiao, C., 2020. Mutations in the pectin methyltransferase QUASIMODO2 influence cellulose biosynthesis and wall integrity in *Arabidopsis*. Plant Cell 32(11), 3576–3597. 10.1105/tpc.20.00252.

ElectronMachine, 2017. Measuring total soluble solids with refractometers. Accessed June 2023. https://blog.electronmachine.com/2017/11/measuring-total-soluble-solids-with.html#:~:text=Total%20Soluble%20Solids%20(TSS)%20refers,of%20citrus%20fruit%20are%20sugars.

El-Taher, A.M., Elzilal, H.A., El-Raouf, H.S., Mady, E., Alshallash, K.S., Alnefaie, R.M., Mahdy, E.M.B., Ragab, O.G., Emam, E.A., Alaraidh, I.A., Randhir, T.O., Ibrahim, M.F.M., 2023. Characterization of some *Cichorium* Taxa grown under Mediterranean climate using morphological traits and molecular markers. Plants 12(2), 388. 10.3390/plants12020388.

Emery, N.J., Offord, C.A., 2019. Environmental factors influencing fruit production and seed biology of the critically endangered *Persoonia pauciflora* (Proteaceae). Folia Geobot 54, 99–113. 10.1007/s12224-019-09343-6.

Etienne, A., Genard, M., Lobit, P., Mbeguie-A-Mbeguie, D., Bugaud, C., 2013. What controls fleshy fruit acidity? A review of malate and citrate accumulation in fruit cells. J Exp Bot 64(6), 1451–1469. 10.1093/jxb/ert035.

Garcia-Munoz, M.C., Henao-Rojas, J.C., Moreno-Rodriguez, J.M., Botina-Azain, B.L., Romero-Barrera, Y., 2021. Effect of rootstock and environmental factors on fruit quality of Persian lime (Citrus latifolia Tanaka) grown in tropical regions. 103, 104081. 10.1016/j.jfca.2021.104081.

Gnan, S., Priest, A., Kover, P.X., 2014. The genetic basis of natural variation in seed size and seed number and their trade-off using *Arabidopsis thaliana* MAGIC lines. Genetics 198(4), 1751–1758. 10.1534/genetics.114.170746.

Gogoi, M., 2021. Assam Lemon: a magical fruit. BioNE 25, 3790.

Government of Assam Database. 2022. Area, production and average yield of some major horticultural crops of Assam. Accessed August 2023. https://des.assam.gov.in/sites/default/files/swf_utility_folder/departments/ecostat_medhassu_in_oid_3/menu/document/statistical_handbook_all_pages_final_20-03-2023_without_price.pdf.

Gusakov, A.V., Kondratyeva, E.G., Sinitsyn, A.P., 2011. Comparison of two methods for assaying reducing sugars in the determination of carbohydrase activities. Int J Anal Chem 2011, 1–4. 10.1155/2011/283658.

Hore, D.K., Barua, U., 2004. Status of Citriculture in North Eastern region of India - a review. Agric Rev 25(1), 1–15.

Hsouna, A.B., Sadaka, C., Mekinic, I.G., Garzoli, S., Svarc-Gajic, J., Rodrigues, F., Morais, S., Moreira, M.M., Ferreira, E., Spigno, G., Brezo-Borjan, T., Akacha, B.B., Saad, R.B., Delerue-Matos, C., Mnif, W., 2023. The chemical variability, nutraceutical value, and food-industry and cosmetic applications of Citrus plants: a critical review. Antioxidants 12(12), 481. 10.3390/antiox12020481.

Huang, X.Y., Wang, C.K., Zhao, Y.W., Sun, C.H., Hu, D.G., 2021. Mechanisms and regulation of organic acid accumulation in plant vacuoles. Hortic Res 8, 227. 10.1038/s41438-021-00702-z.

Jain, J.R., Timsina, B., Satyan, K.B., Manohar, S.H., 2017. A comparative assessment of morphological and molecular diversity among *Sechium edule* (Jacq.) Sw. accessions in India. 3 Biotech 7, 106. 10.1007/s13205-017-0726-5.

Janeczko, D.B., Timmons, M.B., 2019. Effects of seeding pattern and cultivar on productivity of Baby Spinach (*Spinacia oleracea*) grown hydroponically in deep-water culture. Hortic 5(1): 20. 10.3390/horticulturae5010020.

Jing, C., Feng, D., Zhao, Z., Wu, X., Chen, X., 2020. Effect of environmental factors on skin pigmentation and taste in three apple cultivars. Acta Physiol Plant 42, 69. 10.1007/s11738-020-03039-7.

Kashyap, K., Kashyap, D., Nitin, M., Ramchiary, N., Banu, S., 2020. Characterizing the nutrient composition, physiological maturity, and effect of cold storage in Khasi Mandarin (*Citrus reticulata* Blanco). Int J Fruit Sci 20(3), 521–540. 10.1080/15538362.2019.1666334.

Khalil, H.P.S.A., Hossain, M.S., Rosamah, E., Azli, N.A., Saddon, N., Davoudpoura, Y., Islam, M.N., Dungani, R., 2015. The role of soil proper’ w: w R w Sustainable Energy Rev 43, 1006–1015. 10.1016/j.rser.2014.11.099.

Khamsucharit, P., Laohaphatanalert, K., Gavinlertvatana, P., Sriroth, K., Sangseethong, K., 2018. Characterization of pectin extracted from banana peels of different varieties. Food Sci Biotechnol 27(3), 623–629. 10.1007%2Fs10068-017-0302-0.

Khatiwora, E., Adsul, V.B., Torane, R.C., Gaikwad, S., Deshpande, N.R., Kashalkar, R.V., 2017. Spectroscopic determination of total phenol and flavonoid contents of *Citrus limon* peel from North Eastern region of India. J Drug Deliv Ther 7(1), 21–24. 10.22270/jddt.v7i1.1368.

Kumar, V., Sharma, V., Singh, L., 2018. Pectin from fruit peels and its uses as pharmaceutical and food grade: a descriptive review. European J Biomed Pharm Sci 5(5), 185–189.

Lado, J., Alos, E., Manzi, M., Cronje, P.J.R., Gomez-Cadenas, A., Rodrigo, M.J., Zacarias, L., 2019. Light regulation of carotenoid biosynthesis in the peel of Mandarin and Sweet Orange fruits. Front Plant Sci 10, 1288. 10.3389/fpls.2019.01288.

Li, N., Wang, J., Wang, B., Huang, S., Hu, J., Yang, T., Asmutola, P., Lan, H., Qinghui, Y., 2021. Identification of the carbohydrate and organic acid metabolism genes responsible for Brix in tomato fruit by transcriptome and metabolome analysis. Front Genet 12, 714942. 10.3389/fgene.2021.714942.

Liao, L., Dong, T., Qiu, X., Rong, Y., Wang, Z., Zhu, J., 2019. Nitrogen nutrition is a key modulator of the sugar and organic acid content in Citrus fruit. PLoS ONE 14(10), e0223356. 10.1371/journal.pone.0223356.

Liliane, T.N., Charles, M.S., 2019. Factors affecting yield of crops, in: Amanullah, (Ed.), Agronomy. IntechOpen., London, pp. 1–16. 10.5772/intechopen.90672.

Morgan, J.B., Connolly, E.L., 2013. Plant-soil interactions: nutrient uptake. Nat Educ knwl 4(8), 2.

Mukhim, C., Nath, A., Deka, B., Swer, T.L., 2015. Changes in physico-chemical properties of Assam lemon (*Citrus limon* burm.) at different stages of fruit growth and development. Bioscan 10(2), 535–537.

Neri, J., Wendt, T., Silva, CP., (2018). Natural hybridization and genetic and morphological variation between two epiphytic bromeliads. AoB Plants 10(1): plx061. 10.1093/aobpla/plx061.

Pacheco-Hernandez, Y., Villa-Ruano, N., Lozoya-Gloria, E., Barrales-Cortes, C.A., Jimenez-Montejo, F.E., Cruz-Lopez, M.C., 2021. Influence of environmental factors on the genetic and chemical diversity of *Brickellia veronicifolia* populations growing in fragmented Shrublands from Mexico. Plants 10(2), 325. 10.3390/plants10020325.

Padayatty, S.J., Levine, M., 2016. Vitamin C physiology: the known and the unknown and Goldilocks. Oral Dis 22(6), 463–493. 10.1111/odi.12446.

Perez-Nicolas, M., Colinas-Leon, T., Alia-Tejacal, I., Pena-Ortega, G., Gonzalez-Andres, F., Beltran- Rodriguez, L., 2021. Morphological Variation in Scarlet Plume (*Euphorbia fulgen*s Karw ex Klotzsch, Euphorbiaceae), an Underutilized Ornamental Resource of Mexico with Global Importance. Plants 10(10), 2020. 10.3390/plants10102020.

Porter, D.R., Bisson, C.S., Allinger, H.W., 1940. Factors affecting the total soluble solids, reducing sugars, and sucrose in watermelons. Hilgardia 13(2), 31–66.

Premachandran, A., Dhayasree, K., Kurein, S., 2019. Seedless fruits: fruits of future. J Pharmacogn Phytochem 8(6), 1053–1059.

Qi, J., Liu, W., Jiao, T., Hamblin, A., 2020. Variation in morphological and physiological characteristics of wild *Elymus nutans* ecotypes from different altitudes in the Northeastern Tibetan Plateau. Sens App Agric Environ Monit 2020, 2869030. 10.1155/2020/2869030.

Reddy, A., Norris, D.F., Momeni, S.S., 2016. The pH of beverages in the United States. J Am Dent Assoc 147(4), 255–263. 10.1016/j.adaj.2015.10.019.

Saeid, A., Ahmed, M., 2021. Citrus fruits: nutritive value and value-added products, in: Khan, M.S., Khan, I.A. (Eds.), Citrus. IntechOpen., London, pp. 1–18. 10.5772/intechopen.95881.

Samtiya, M., Aluko, R.E., Dhewa, T., Moreno-Rojas, J.M., 2021. Potential health benefits of plant food- derived bioactive components: an overview. Foods 10(4), 839. 10.3390/foods10040839.

Satpathy, L., Pradhan, N., Dash, D., Baral, P.P., Parida, S.P., 2021. Quantitative determination of Vitamin C concentration of common edible food sources by redox titration using iodine solution. Lett Appl Nano Bio Sci 10(3), 2361–2369. 10.33263/LIANBS103.23612369.

Schjoerring, J.K., Cakmak, I., White, P.J., 2019. Plant nutrition and soil fertility: synergies for acquiring global green growth and sustainable development. Plant Soil 434, 1–6. 10.1007/s11104-018-03898-7.

Sheikh, K.H.A., Singh, B., Haokip, S.W., Shankar, K., Debbarma, R., Devi, A.G., Nengparmoi, T.H., 2021. Response of yield and fruit quality to foliar application of micronutrients in Lemon [*Citrus limon* (L.) Burm.] cv. Assam Lemon. J Hortic 16(2), 144–153.

Silver, W.L., Perez, T., Mayer, A., Jones, A.R., 2021. The role of soil in the contribution of food and feed. Trans R Soc B 376, 20200181. 10.1098/rstb.2020.0181.

Smith, M.R., Hodecker, B.E.R., Fuentes, D., Merchant, A., 2022. Investigating nutrient supply effects on plant growth and seed nutrient content in common bean. Plants 11(6), 737. 10.3390/plants11060737.

Sreekumar, V.B., Renuka, C., Suma, T.B., Balasundaran, M., 2006. Taxonomic reconsideration of *Calamus rivalis* Thw. ex Trim. and *C. metzianus* Schlecht (Arecaceae) through morphometric and molecular analyses. Bot Stud 47, 443–452.

Srivastava, A.K., Singh, S., 2007. Analysis of Citrus orchard efficiency in relation to soil properties. J Plant Nutr 30(12), 2077–2090. 10.1080/01904160701700566.

Susandarini, R., Subandiyah, S., Rugayah., Daryono, B.S., Nugroho, L.H., 2013. Assessment of taxonomic affinity of indonesian Pummelo (*Citrus maxima* (Burm.) Merr.) based on morphological characters. American J Agric Biol Sci 8(3), 182–190. 10.3844/ajabssp.2013.182.190.

Tasisa, J., Mohammed, W., Hussien, S., Kumar, V., 2018. Genetic control of inheritance of fruit quality attributes in tomato (*Solanum lycopersicum*). Agric Res 7, 120–128. 10.1007/s40003-018-0314-x.

Thakur, S., 2010. World’s first experimental plantation of grafted Assam lemon raised at Kahikuchi. Assam Tribune. Accessed on 1/2/2023. https://assamtribune.com/worlds-first-experimental-plantation-of-grafted-assam-lemon-raised-at-kahikuchi#:~:text=Using%20the%20seeds%20of%20China,others%20and%20had%20seedless%20fruit.

TNAU Agritech Portal. 2013. Resource management: soil: soil sampling procedure. Accessed on 12/8/2023. https://agritech.tnau.ac.in/agriculture/agri_soil_sampling.html#:~:text=For%20shallow%20rooted%20crops%2C%20collect,up%20to%2030%20cm%20depth.

Tosso, F., Doucet, J.L., Dainou K., Fayolle, A., Hambuckers, A., Doumenge, C., Agbazahou, H., Stoffelen, P., Hardy, O.J., 2019. Highlighting convergent evolution in morphological traits in response to climatic gradient in African tropical tree species: The case of genus *Guibourtia* Benn. Ecol Evol 9(23), 13114–13126. 10.1002/ece3.5740.

Uresti-Porras, J., Fuente, M.C.F., Benavides-Mendoza, A., Olivares-Saenz, E., Cabrera, R.I., Juarez- Maldonado, A., 2021. Effect of graft and nano ZnO on nutraceutical and mineral content in ball paper. Plants 10(12), 1–18. 10.3390%2Fplants10122793.s

Vanitha, T., Khan, M., 2019. Role of pectin in food processing and food packaging, in: Masuelli, M.A. (Ed.), Pectins, IntechOpen., London, pp 1–21. 10.5772/intechopen.83677.

Velasco, R., Licciardello, C., 2014. A genealogy of the *Citrus* family. Nat Biotechnol 32: 640–642. 10.1038/nbt.2954.

Venkatanagaraju, E., Bharathi, N., Sindhuja, R.H., Chowdhury, R.R., Sreelekha, Y., 2019. Extraction and purification of pectin from agro-industrial wastes, in: Masuelli, M. (Ed.), Pectins., London, pp. 1–16. 10.5772/intechopen.85585.

Vita, F., Franchina, F.A., Taiti, C., Locato, V., Pennazza, G., Santonico, M., Purcaro, G., Gara, L.D., Mancuso, S., Mondello, L., Alpi, A., 2018. Environmental conditions influence the biochemical properties of the fruiting bodies of *Tuber magnatum* Pico. Sci Rep 8, 7243. 10.1038/s41598-018-25520-7.

Wang, L., Li, J., Zhao, J., He, C., 2015. Evolutionary developmental genetics of fruit morphological variation within the Solanaceae. Front Plant Sci 6, 248. 10.3389/fpls.2015.00248.

Wang, M., Zhang, J., Guo, Z., Guan, Y., Qu, G., Liu, J., Guo, Y., Yan, X., 2020. Morphological variation in *Cynodon dactylon* (L.) Pers., and its relationship with the environment along a longitudinal gradient. Hereditas 157, 4. 10.1186/s41065-020-00117-1.

Wu, G.A., Prochnik, S., Jenkins, J., Salse, J., Hellsten, U., Murat, F., Perrier, X., Ruiz, M., Scalabrin, S., Terol, J., Takita, M.A., Labadie, K., Poulain, J., Couloux, A., Jabbari, K., Cattonaro, F., Fabbro C.D., Pinosio, S., Zuccolo, A., Chapman, J., Grimwood, J., Tadeo, F.R., Estornell, L.H., Munoz-Sanz, J.V., Ibanez, V., Herrero- Ortega, A., Aleza, P., Perez-Perez, J., Ramon, D., Brunel, D., Luro, F., Chen, C., Farmerie, W.G., Desany, B., Kodira, C., Mohiuddin, M., Harkins, T., Fredrikson, K., Burns, P., Lomsadze, A., Borodovsky, M., Reforgiato, G., Freitas-Astua, J., Quetier, F., Navarro, L., Roose, M., Wincker, P., Schmutz, J., Morgante, M., Machado, M.A., Talon, M., Jaillon, O., Ollitrault, P., Gmitter, F., Rokhsar, D., 2014. Sequencing of diverse Mandarin, Pummelo and Orange genomes reveals complex history of admixture during citrus domestication. Nat Biotechnol 32(7): 656–62. 10.1038/nbt.2906.

Ye, X., 2018. Phytochemicals in Citrus: applications in functional foods, first ed. CRC Press, Florida. 10.4324/9781315369068.

Zhang, T., Hong, Y., Zhang, X., Yuan, X., Chen, S., 2022. Relationship between key environmental factors and the architecture of fruit shape and size in near-isogenic lines of cucumber (*Cucumis sativus* L.). Int J Mol Sci 23(22), 14033. 10.3390/ijms232214033.

